# The evolution of an RNA-based memory of self in the face of genomic conflict

**DOI:** 10.1101/2022.06.17.496645

**Authors:** Pinelopi Pliota, Hana Marvanova, Alevtina Koreshova, Yotam Kaufman, Polina Tikanova, Daniel Krogull, Andreas Hagmüller, Sonya A. Widen, Dominik Handler, Joseph Gokcezade, Peter Duchek, Julius Brennecke, Eyal Ben-David, Alejandro Burga

## Abstract

Distinguishing endogenous genes from selfish ones is essential for germline integrity. In animals, small regulatory RNAs play a central role in this process; however, the underlying principles are largely unknown. To fill this gap, we studied how selfish toxin-antidote elements (TAs) evade silencing in the nematode *Caenorhabditis tropicalis*. We found that the *slow-1/grow-1* TA is active only when maternally inherited. Surprisingly, this parent-of-origin effect stems from a regulatory role of the toxin’s mRNA: maternal *slow-1* mRNA—but not SLOW-1 protein—licenses *slow-1* expression in the zygote by counteracting piRNAs. Our results indicate that epigenetic licensing— known to play a role in *C. elegans* sex-determination—is likely a common mechanism that hinders the spread of selfish genes in wild populations while ensuring a lasting memory of self in the germline.

## Main Text

The germline is unique in that it is the only immortal lineage of animals. This feature makes it an attractive target for parasitic elements and viruses seeking to perpetuate themselves at the expense of their host. In the face of this existential threat, organisms have evolved a variety of mechanisms to defend their genomes, all of which have one thing in common: to neutralize the intruder, they must first distinguish endogenous from foreign nucleic acids. To put it another way, these molecular circuits have evolved the ability to distinguish between “self” and “non-self”. Failure to do so can have drastic consequences for the germline, such as the reactivation of transposons or the silencing of endogenous genes, both of which result in sterility [1–4]. Nucleic acid complementary base-pairing via small regulatory RNAs (sRNAs, <30 bp in length) bound to Argonaute proteins is a recurrent feature of defense mechanisms across the tree of life [5]. Yet, how organisms employ sRNAs to efficiently distinguish self from non-self and develop a long-lasting memory of endogenous transcripts is still poorly understood.

Originally discovered in *Drosophila*, PIWI-interacting RNAs (piRNAs) are a class of sRNA that play a central role in silencing transposable elements and selfish repeats in the animal germline [3, 6]. In line with this view, *C. elegans* piRNAs—also known as 21U-RNAs—can efficiently silence transgenic reporters made of exogenous sequences [7–10]. To do so, piRNAs first form a complex with the PIWI protein PRG-1 and target complementary sequences, allowing for imperfect base-pairing. This then leads to the production of secondary small RNAs, 22G-RNAs, via the recruitment of RNA-dependent RNA polymerases (RdRPs), which in turn are loaded onto worm-specific Argonaute proteins that induce epigenetic silencing in the nucleus [11]. Although the targeting rules of piRNAs and the components of their pathway have been extensively studied, the physiological role of piRNAs beyond silencing of synthetic transgenes is far less understood. Counterintuitively, most *C. elegans* piRNAs do not target transposable or viral elements but instead are complementary to thousands of endogenous genes [12]. This observation raises a critical question—how are endogenous genes not silenced in the germline? As a counterbalance to the silencing effect of piRNAs, two main mechanisms have been put forward: a periodic 10 bp motif of A_n_/T_n_ clusters (PATCs), and the CSR-1 Argonaute [13–16]. PATCs are typically found in the introns of germline-expressed genes that reside in repressive chromatin and can promote the expression of transgenes in the germline [14, 17]. Unlike all other Argonautes, CSR-1 can activate silenced transgenes via ‘RNAa’ (RNA-induced epigenetic gene activation) and this activity has been proposed to rely also on 22G-RNAs [15]. However, whether PATCs or CSR-1 protect endogenous genes from silencing in the germline is still the subject of active research.

Altogether, multiple lines of evidence suggest that organisms discriminate self from non-self via a delicate and ever-changing balance between interacting Argonaute proteins and their associated sRNAs, as well as their local chromatin environment. In this manuscript, instead of relying on synthetic transgenes, we leverage the unique biology of toxin-antidote elements (TAs)—selfish genes that spread in the wild by exploiting maternal-zygotic interactions—to study this process [18]. Unlike transposons and viruses, many TAs originate from endogenous genes via gene duplication, and thus provide a unique opportunity to study how the memory of gene expression is established and evolves in the face of genomic conflict, as genes transition from friend to foe.

### The *slow-1/grow-1* selfish TA has a parent-of-origin effect

*Caenorhabditis tropicalis* is a hermaphroditic nematode that—unlike its cosmopolitan relative *C. elegans*—inhabits exclusively equatorial regions [19]. While studying genetic incompatibilities between two *C. tropicalis* wild isolates NIC203 (Guadeloupe, France) and EG6180 (Puerto Rico, USA), we discovered a novel maternal-effect TA, which we named *slow-1/grow-1* [20]. This selfish element is located in NIC203 Chr. III and comprises three tightly linked genes: a maternally expressed toxin, *slow-1*, and two identical and redundant antidotes, *grow-1.1* and *grow-1.2*, which are expressed zygotically. For simplicity, we will collectively refer to the two antidotes as *grow-1* unless specifically noted (Figure S1A; see discussion in Supplementary text). In crosses between TA carrier and non-carrier strains, heterozygous mothers poison all their eggs but only progeny that inherit the TA can counteract the toxin by zygotically expressing its antidote. Consequently, homozygous non-carriers are exclusively poisoned by the toxin (Figure 1A). While wild-type worms typically take two days to develop from the L1 stage to the onset of egg laying, embryos poisoned by SLOW-1 take on average four days, and some even longer. This delay imposes a high fitness cost and favors the spread of the selfish element in the population.

**Figure 1.**
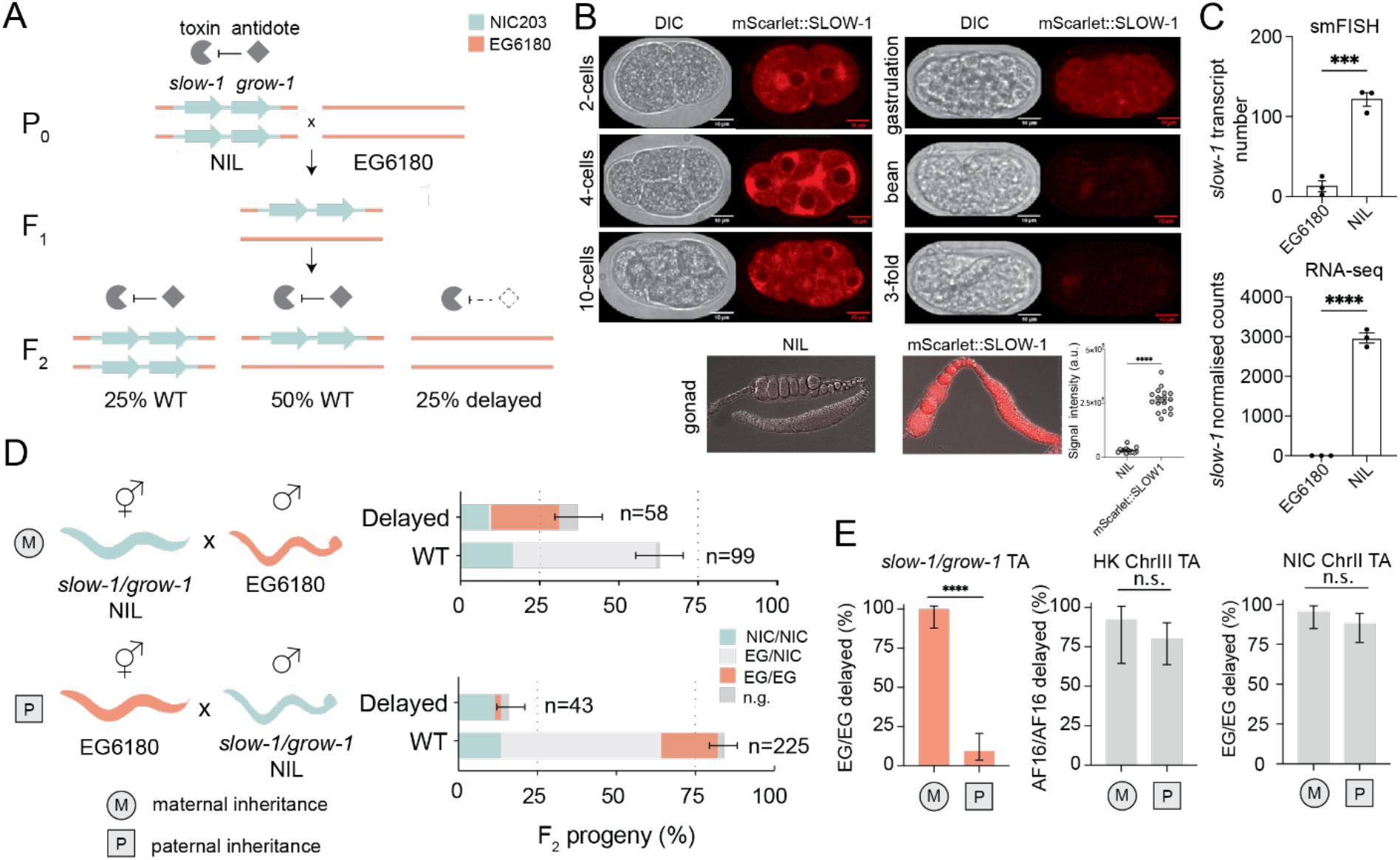
*slow-1/grow-1*, a selfish element with a parent-of-origin effect. **(A)** Model illustrating the mechanism of action of the *slow-1/grow-1* TA. In crosses between the carrier strain (slow-1/grow-1 Chr. III NIL) and non-carrier strain (EG6180), 25% of the F_2_ progeny is developmentally delayed because EG/EG homozygotes did not inherit the TA and cannot express the antidote to counteract the maternally deposited toxin. **(B)** Expression pattern of mScarlet::SLOW-1 during embryonic development (top) and hermaphroditic gonad (bottom). Quantification of signal intensity in gonads of NIL and mScarlet::SLOW-1 strains (t-test, p<0.0001). **(C)** Quantification of *slow-1* mRNA expression by smFISH (N=3, n=16-30 per repeat, unpaired t-test, p=0.0006) and RNA-seq in both NIL and EG6180 parental strains (unpaired t-test, p<0.0001). **(D)** Reciprocal crosses between the *slow-1/grow-1* TA NIL (cyan) and the EG6180 parental strain (orange). Maternal or paternal inheritance refers to the *slow-1/grow-1* locus. Worms with a significant developmental delay or laval arrest were categorized as delayed, otherwise classified as wild type (WT). Sample sizes are shown for each phenotypic class (n). Error bars indicate 95% binomial confidence intervals calculated with the Agresti-Coull method. Not genotyped because of technical problems (n.g.). Each cross was independently performed at least twice with identical results (see Excel Table1 for raw data). **(E)** Activity of various TA elements in reciprocal crosses. Penetrance of the toxin, the percentage of F_2_ non-carrier individuals that are phenotypically affected, is used as a proxy for TA activity. (*slow-1/grow-1* TA: n_M_=34, n_P_=53, p<0.0001, HK ChrIII TA: n_M_=13, n_P_=35, p=0.42 and NIC ChrII TA: n_M_=44, n_P_=50, p=0.27, respectively; Fisher’s exact test; *p ≤ 0.05, ** p ≤ 0.01, *** p ≤ 0.001, **** p ≤ 0.0001)

To confirm the maternal deposition of SLOW-1 protein, we tagged the endogenous *slow-1* locus with an N-terminal mScarlet fluorescent tag. The resulting fusion toxin was fully active (Figure S2A). In agreement with its maternal-effect, we found that SLOW-1 was present in the gonads of hermaphrodites throughout their life cycle and loaded into eggs prior to fertilization (Figure 1B). In early embryos, SLOW-1 appeared to be associated with the nuclear envelope and was quickly degraded during embryogenesis. SLOW-1 was not detectable in the soma by the time embryos reached the comma-stage. This fast clearance likely reflects the activity of the GROW-1 antidote. To better study the inheritance of *slow-1/grow-1* TA, we previously generated a Near-Isogenic Line strain containing the *slow-1/grow-1* NIC203 Chr. III locus in an otherwise EG6180 background, hereinafter “NIL” [20]. As expected, *slow-1* mRNA was detected in the NIL but not in EG6180 (Figure 1C). As previously reported, in crosses between NIL hermaphrodites and EG6180 males, the toxin induced developmental delay in all the F_2_ homozygous non-carrier (EG/EG) individuals (100% delay; n=34; Figure 1D and 1E). However, we noticed an unexpected pattern of inheritance when performing the reciprocal cross. If EG6180 hermaphrodites were mated to *slow-1/grow-1* NIL males, most of their F_2_ EG/EG progeny were not developmentally delayed but phenotypically wild-type (9.4% delay; n=53; *p* ≤ 0.0001) (Figure 1D and 1E). This was surprising because previously known TAs—including the *peel-1/zeel-1* and *sup-35/pha-1* loci in *C. elegans*, and the *Medea* locus in *Tribolium*—affect non-carrier individuals regardless of whether the element is inherited from the maternal or paternal lineage (Figure S3) [21–23].

We also investigated the inheritance pattern of two TAs that we recently discovered in *C. tropicalis* and *C. briggsae* that cause developmental delay. However, we found no evidence of a parent-of-origin effect, indicating that this is not a general feature of non-lethal toxins (Figure 1E). Mito-nuclear incompatibilities could not explain the observed pattern because both parental lines, the NIL and EG6180, carry the same mito-genotype (Figure S4). Moreover, *C. tropicalis*, like all nematodes of the Rhabditida group, lacks *de no*vo methyltransferases, making the involvement of mammalian-like epigenetic imprinting an unlikely scenario [24]. Because parent-of-origin effects are extremely rare in nematodes and all reported cases involve transgenic reporters, we set out to investigate *slow-1/grow-1* further [25, 26].

### A related *slow-2/grow-2* selfish TA has no parent-of-origin effect

The presence of the *slow-1/grow-1* TA in the NIL explains why F_2_ non-carrier progeny (EG/EG) are delayed (Figure 1A). However, it does not account for the fact that ∼1/3 of F_2_ homozygous TA carriers (NIC/NIC) are also delayed in crosses between the NIL and EG6180 (Figure 1D). We previously hypothesized that this inheritance pattern could emerge from an independent incompatibility factor present in EG6180 within the homologous boundaries of the NIC203 introgression [20]. Since genetic interactions between *slow-1/grow-1* and the putative EG6180 element could influence the activity of the former and perhaps cause the parent-of-origin effect, we set out to map and characterize the latter.

We first sequenced and assembled the EG6180 genome using a combination of Nanopore long-reads and Illumina short-reads and annotated the genome using RNA-seq data (see methods). We then searched for genes that were either missing, mutated or highly divergent within the NIC203 40kb introgression region but present in EG6180. We found a pair of highly divergent but homologous genes to the *slow-1/grow-1* TA in tight genetic linkage (Figure 2A). We named the new element *slow-2/grow-2*. *slow-2* transcripts were readily detectable in EG6180 but absent in the NIL (Figure 2B). Therefore, we concluded that EG6180 encodes *slow-2/grow-2* at the same locus at which NIC203 encodes *slow-1/grow-1*, and the NIL is equivalent to EG6180 with *slow-2/grow-2* replaced by *slow-1/grow-1*.

**Figure 2.**
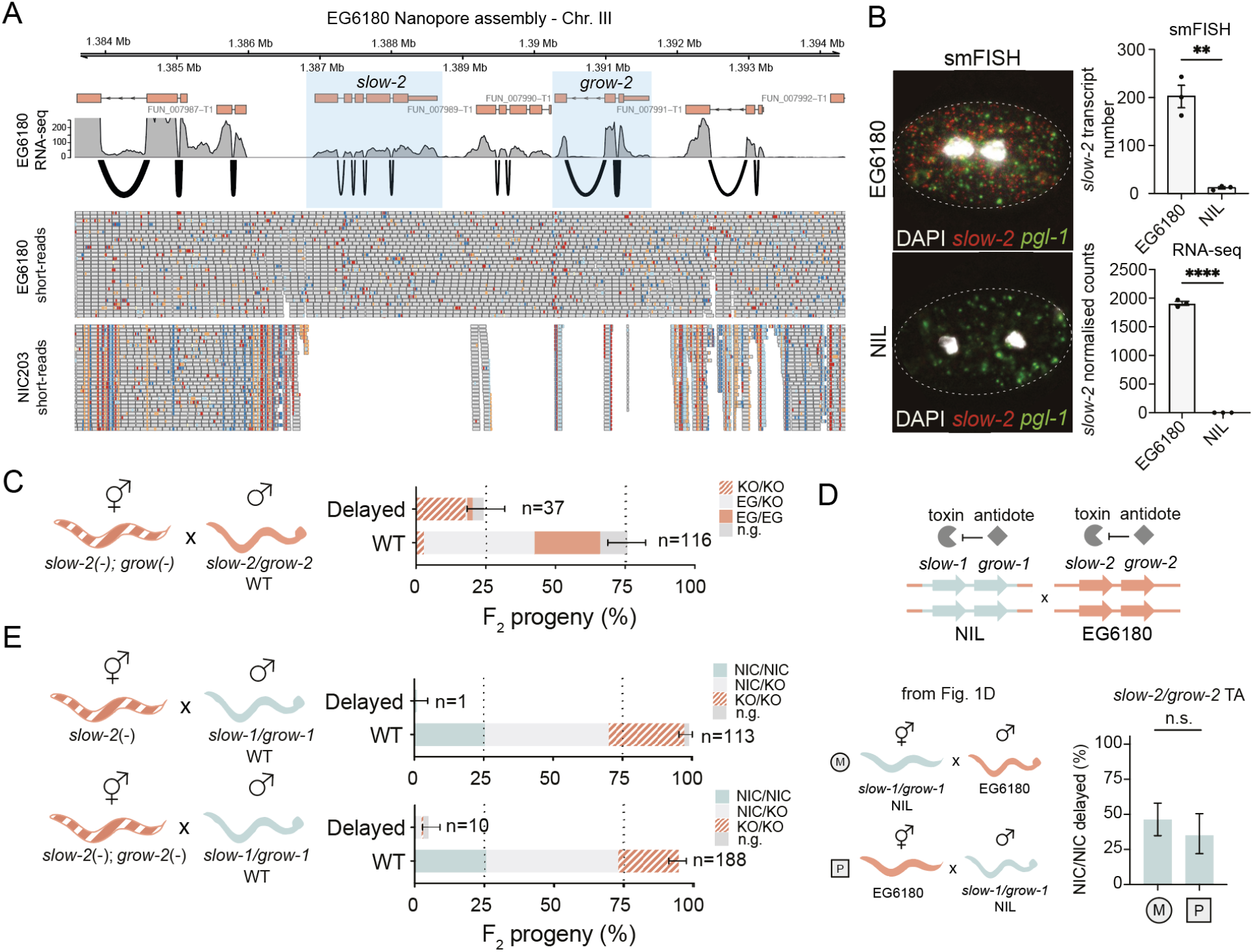
Conflict between selfish TAs leads to evolutionary diversification. **(A)** Alignment of Illumina short-reads to the EG6180 *de novo* assembly. Highlighted region is Chr. III EG6180 region homologous to NIC203 *slow-1/grow-1* **(B)** *slow-2* expression in EG6180 2-cell stage embryos by smFISH (left and top right, N=3, n=16-30 per repeat, unpaired t-test, p=0.0013). *pgl-1* serves as a positive control. *slow-2* expression in EG6180 and absence in the NIL was also confirmed by RNA-seq data (bottom right, n=3). **(C)** Mating of *slow-2(-) grow-2(-)* double mutant NIL (hermaphrodites) to the parental EG6180 line causes developmental delay of homozygous double-mutant F_2_ individuals indicating that *slow-2/grow-2* is a TA (see legend of Figure 1 for explanation of symbols and statistics). **(D)** Two TAs are active in crosses between the NIL and EG6180: *slow-1/grow-1* and *slow-2/grow-2*, respectively (Figure 1D and 1E). The penetrance of the *slow-2/grow-2* TA is incomplete; however, the toxin is equally active when maternally or paternally inherited (n_M_=40, n_P_=67, Fisher’s exact test, p=0.31). **(E)** Genetic crosses indicate that the parent-of-origin effect in *slow-1* is independent of *slow-2* and *slow-2/grow-2* activity.

To test whether the *slow-2/grow-2* locus is a *bona fide* TA, we generated a *slow-2(-)*/*grow-2(-)* double mutant strain using CRISPR/Cas9 in the parental EG6180 background. Both mutant alleles introduce a premature stop codon that is predicted to generate a null allele (Figure S5). We then crossed *slow-2(-)/grow-2(-)* double mutant hermaphrodites to WT EG6180 males and inspected their F_2_ progeny (Figure 2C). If these two genes encode a toxin and an antidote, we would expect the double mutant to behave like a NIC203 susceptible allele [20]. Indeed, we observed 23.1% delayed worms among the F_2_ progeny (n=160), while 87% (n=31) of *slow-2(-)/grow-2(-)* double mutant individuals were developmentally delayed (Figure 2C), in agreement with the TA model. As a control, we observed only background levels of developmental defects in the *slow-2(-)* single mutant and *slow-2(-)/grow-2(-)* double mutant parental lines (3.8%; n=130 and 0.72%; n=139, respectively). Thus, both *slow-2* and *grow-2* are dispensable for the normal development of *C. tropicalis*, in line with their selfish role.

To determine whether the toxin was maternally or paternally transmitted to the progeny, we performed backcrosses of heterozygous F_1_ hermaphrodites or males to the parental *slow-1/grow-1* NIL strain. We observed 29.4% (n=139) delayed individuals among the F_2_ progeny from a cross between F_1_ hermaphrodites and NIL males but only background levels of defects when F_1_ males were crossed to NIL hermaphrodites (3%, n=100), indicating that *slow-2* is a maternal-effect toxin (Figure S6). Together, these results show that *slow-2/grow-2* comprises a novel maternal-effect TA, which delays the development of worms independently of *slow-1/grow-1*, but unlike s*low-1/grow-1* does not show a parent-of-origin effect (Figure 2D). *slow-1/grow-1* and *slow-2/grow-2* share a common evolutionary origin but have diverged to a degree that they are effectively in direct genetic conflict. In crosses between NIC203 and EG6180, each TA selectively targets individuals homozygous for the other element (Figure 1D and Figure S7).

The SLOW-1 and SLOW-2 toxins share 32.6% identity at the protein level, whereas their respective antidotes, GROW-1 and GROW-2, are more conserved—71.0% identical (Figure S7). Despite their relatively low identity, SLOW-1 and SLOW-2 are predicted to have a very similar domain architecture. Both toxins are homologous to nuclear hormone receptors (NHR), a large family of ligand-regulated transcription factors [27]. Like all NHRs, both toxins contain a ligand binding domain but unlike NHRs, they lack a DNA binding domain. Instead, they contain two predicted transmembrane domains on their C-termini (Figure S7). Since NHRs are known to form heterodimers, we tested whether the *slow-1/grow-1* parent-of-origin effect was dependent on the activity of SLOW-2. To do so, we crossed either *slow-2(-)* or *slow-2(-)/grow-2(-)* double knockout EG6180 hermaphrodites to *slow-1/grow-1* NIL males and inspected their F_2_ progeny (Figure 2E). As expected from the absence of the SLOW-2 toxin, the vast majority of F_2_ NIC/NIC individuals were WT in both crosses (96,6%, n=30 and 100%, n=51 respectively, Figure 2E). In addition, virtually all F_2_ EG/EG progeny were also wild-type (0% delay; n=31 and 2.2% delay; n=44; Figure 2E), indicating that the *slow-1/grow-1* parent-of-origin effect is independent of *slow-2/grow-2* activity.

### The *slow-1* maternal toxin is epigenetically licensed

To explore the contribution of *slow-1* to the parent-of-origin effect, we performed reciprocal crosses between the wild-type NIL and a *slow-1(-)/grow-1(-)* double mutant NIL strain, where both the toxin and antidote carry null frameshift mutations (Figure 3A). Analogous to crosses between NIL hermaphrodites and EG6180 males (Figure 1D), when wild type NIL hermaphrodites were mated to the double mutant males, 28.9% (n=190) of the F_2_ progeny were delayed while all the homozygous double mutant individuals were delayed (100%, n=31, Figure 3A). However, much to our surprise, in the reciprocal cross, we observed the same inheritance pattern. Overall, 22.1% (n=140) of the F_2_ progeny were delayed and delayed individuals were homozygous double mutants (95.8%; n=24), indicating that *slow-1/grow-1* was fully active when inherited via the paternal lineage (Figure 3A). Our results thus indicated that the *slow-1/grow-1* double mutant and EG6180 haplotypes were not equivalent with regard to their ability to license a paternal copy of *slow-1/grow-1*. Having already ruled out a direct contribution of *slow-2/grow-2* to the parent-of-origin effect, we reasoned that the most parsimonious explanation for this observation is that expression of the maternally inherited slow*-1* allele was necessary to license the paternal one. Furthermore, given that *slow-1* was able to license a paternal allele despite carrying a frameshift null mutation, we hypothesized that *slow-1* mRNA—but not SLOW-1 protein—is necessary for licensing.

**Figure 3.**
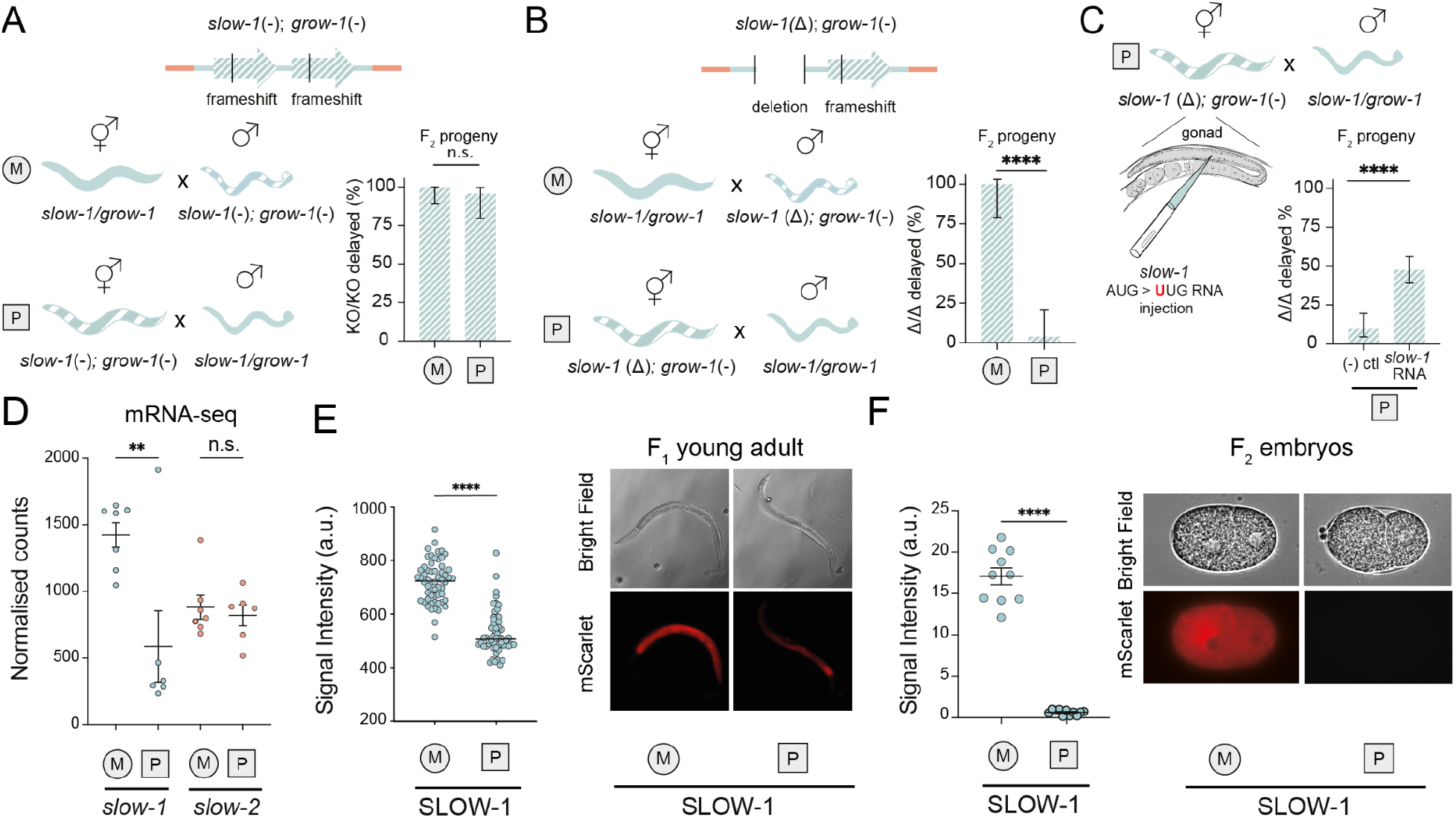
Maternal *slow-1* transcripts are necessary and sufficient to license the expression of zygotic *slow-1*. **(A)** In the *slow-1(-)/grow-1(-)* double mutant NIL strain, both *slow-1* and *grow-1* carry frameshift mutations (top). Reciprocal crosses between the NIL and *slow-1(-)/grow-1(-)* double mutants indicate that the TA is fully active both when maternally and paternally inherited (bottom, n_M_=31, n_P_=24, Fisher’s exact test p=0.43) **(B)** In the *slow-1(Δ)/grow-1(-)* mutant NIL strain, the coding region of *slow-1* has been deleted and *grow-1* carries the frameshift mutation as in (A) (top). Reciprocal crosses between the NIL and *slow-1(Δ)/grow-1(-)* mutants indicate that *slow-1/grow-1* is only active when maternally inherited (bottom, n_M_=18, n_P_=25, Fisher’s exact test p<0.0001) **(C)** *In vitro* transcribed *slow-1* RNA with a mutated start codon was injected in the gonad of *slow-1(Δ)/grow-1(-)* double mutant NILs and later crossed to NIL males. Approximately half of their *Δ*/*Δ* F_2_ progeny were delayed. Control mothers were injected with DEPC H_2_O (n_slow-1 RNA_=128, n_ctl_=62, Fisher’s exact test p<0.0001). **(D)** Reciprocal crosses between the NIL and EG6180 followed by RNA-seq of their F_1_ progeny indicates that *slow-1* transcripts are more abundant when maternally inherited. In contrast, *slow-2* transcripts are equally abundant (two-way ANOVA, interaction p=0.0144, Holm-Sidak post hoc test, p_slow-1_=0.0011, p_slow-2_=0.77) **(E)** Reciprocal crosses between mScarlet::SLOW-1 NIL and EG6180 strains. Quantification of total body fluorescence of F_1_ young adults includes signal from the germline and also gut autofluorescence. Each dot represents a single individual. Reciprocal crosses were performed twice (unpaired t-test, n_M_=58, n_P_=58, p<0.0001). Representative images of F_1_ young adults from the reciprocal crosses (right). **(F)** Reciprocal crosses between mScarlet::SLOW-1 NIL and EG6180 strains. Quantification of total fluorescence of F_2_ 2-cell stage embryos (unpaired t-test, n_M_=10, n_P_=11, p<0.0001).

To test this hypothesis, we used CRISPR/Cas9 to delete the full coding region of *slow-1* in an otherwise identical genetic background to the double mutant NIL strain carrying *slow-1(-)/grow-1(-)*. In contrast to the frameshift allele, the deletion (*Δ*) completely removes *slow-1* transcripts. Then, we performed reciprocal crosses between the *slow-1(Δ)/grow-1*(-) NIL strain and the wild-type NIL, and inspected their F_2_ progeny (Figure 3B). When NIL hermaphrodites were crossed to *slow-1(Δ)/grow-1*(-) NIL males, we observed 26.8% (n=190) delay among the F_2_ while all genotyped worms homozygous for the mutant allele were delayed (100%, n=18; Figure 3B) In contrast, when *slow-1(Δ)/grow-1*(-) NIL hermaphrodites were crossed to NIL males, we observed baseline delay among their F_2_ progeny (3.8%, n=129) (Figure 3B). Thus, our results indicate that the *slow-1(-)* allele but not *slow-1(Δ)* is able to license a paternally inherited slow*-1/grow-1* TA and identify *slow-1* mRNA as the licensing signal.

To test whether *slow-1* mRNA is sufficient for licensing, we *in vitro* transcribed *slow-1* RNA and injected it into the gonads of 16 *slow-1(Δ)/grow-1*(-) NIL hermaphrodites, mated those to *slow-1/grow-1* NIL males and inspected their F_2_ progeny. Critically, we mutated the start codon of the slow-1 cDNA that served as a transcription template, resulting in RNA that cannot be translated into SLOW-1 protein. Following injection of noncoding *slow-1* RNA into the gonads of *slow-1(Δ)/grow-1*(-) mothers, there was a significant increase in the proportion of delayed individuals among the F_2_ compared to a control injection (13.8% delayed, n=650 and 4% delayed, n=296 respectively, Fisher’s exact test, p<0.0001). Most (84.7%, n=61) of genotyped delayed individuals were homozygous for the *slow-1(Δ)/grow-1*(-) allele, showing that the effect was highly specific. Overall, injection of *slow-1* RNA increased the proportion of delayed F_2_ individuals among double mutants from 9.6% (n=62) in the control cross to 47.6% (n=128) in the treatment (p<0.0001; Figure 3C). These results show that *slow-1* RNA is sufficient to license a paternally inherited slow*-1/grow-1* allele and that this effect does not depend on SLOW-1 protein. The partial rescue of zygotic *slow-1* expression likely reflects technical limitations in the injection protocol, as we observed a wide range of rescue depending on the injected mother (Figure S8).

Next, we tested the effect of maternal *slow-1* dosage on licensing. To do so, we first used CRISPR/Cas9 to delete 620 bp upstream of the *slow-1* coding region in its endogenous locus and then proceeded to knock out the antidote *grow-1*. RNA-seq of the resulting promoter deletion strain revealed a 176-fold decrease in *slow-1* mRNA levels while neighboring genes were unaffected (Figure S9). We observed only limited abnormal phenotypes in the double mutants despite lacking *grow-1* antidote (8.71% delay, n=70), suggesting that the amount of SLOW-1 made in the promoter deletion line was not sufficient to poison embryos as efficiently as the WT allele. However, when we crossed *slow-1(Δprom)/grow-1*(-) hermaphrodites to NIL males, the paternal allele was fully active: 27.7% of F_2_ individuals were delayed (Figure S9). These results indicate that a 176-fold reduction in *slow-1* maternal mRNA abundance abolishes its toxicity but not its licensing activity, suggesting that licensing involves a catalytic step (Figure S9).

Lastly, to check for the specificity of the licensing mechanism, we crossed *slow-1(Δ)/grow-1*(-) NIL hermaphrodites to EG6180 males and checked whether a paternally inherited *slow-2/grow-2* was still active in the absence of maternal *slow-1* mRNA. Inspection of their F_2_ progeny revealed 24.4% (n=90) affected progeny, out of which 86.3% (n=22) were homozygous for the *slow-1* deletion allele (Figure S10). We thus concluded that, despite sharing homology, paternally inherited *slow-2* does not require maternal *slow-1* mRNA to be efficiently expressed and to poison embryos.

### Lack of licensing leads to decreased SLOW-1 dosage

Having established that *slow-1* maternal mRNA is the licensing signal, we set out to understand its downstream molecular consequences. Maternally expressed *slow-1*, which is loaded in eggs prior to fertilization both as mRNA and protein, is responsible for the *slow-1/grow-1* TA delay-associated phenotype. Thus, we reasoned that the parent-of-origin effect could stem from reduced expression of the toxin in the germline of F_1_ heterozygous mothers. To test this idea, we performed reciprocal crosses between EG6180 and the *slow-1/grow-1* NIL strains, followed by RNA extraction from F_1_ heterozygous young adult hermaphrodites, and transcriptome sequencing by RNA-seq. In agreement with our hypothesis, *slow-1* mRNA levels were significantly lower in F_1_ mothers when *slow-1/grow-1* was paternally inherited (2.4-fold decrease, two-way ANOVA, Holm-Sidak post hoc test, p=0.0011; Figure 3D). Importantly, there was no difference in *slow-2* expression levels between the reciprocal crosses, in line with *slow-2/grow-2* being equally active when inherited from either parent (Figure 3D). The *slow-1* parent-of-origin effect was not exclusive to the recombinant NIL strain, as we observed the same difference in *slow-1* gene expression when performing reciprocal crosses between NIC203 and EG6180 parental strains (Figure S11).

To independently validate the parent-of-origin effect, we took advantage of our endogenously tagged mScarlet::SLOW-1 NIL strain (Figure 1B). We performed reciprocal crosses between the mScarlet::SLOW-1 and EG6180 strains and quantified the fluorescence signal in the germline of their F_1_ progeny. In agreement with both our genetic crosses and RNA-seq experiments, when *slow-1/grow-1* was paternally inherited, SLOW-1 protein levels were significantly lower in the germline of F_1_ individuals (unpaired t-test, n_M_=58, n_P_=58, p<0.0001) (Figure 3E), as well as in F_2_ 2-cell stage embryos (unpaired t-test, n_M_=10, n_P_=11, p<0.0001) (Figure 3F). Moreover, the mScarlet::SLOW-1 toxin only caused developmental delay among F_2_ progeny when maternally inherited, indicating that epigenetically licensed genes do not lose this status after the introduction of an exogenous sequence tag (Figure S2B). Overall, our results show that the lack of activity of the paternally inherited *slow-1/grow-1* allele stems from a reduction of *slow-1* mRNA levels in the germline of the F_1_ mother and, consequently, a reduced dosage of the toxin in F_2_ embryos. To test whether the SLOW-1 dosage indeed correlates with the severity of phenotype, we decided to impair the antidote function in the parental NIL strain that has double the amount of *slow-1* mRNA compared to heterozygous worms (Figure S1B). We found that *grow-1.1*(+/-); *grow-1.2*(-/-) worms were viable but failed to retrieve any viable *grow-1.1*(-/-); *grow-1.2*(-/-) individuals among their progeny. They arrested as larvae (Figure S1B). This result indicates that doubling the dosage of *slow-1* does not cause developmental delay but is instead lethal. Collectively, our results show that the maternal *slow-1* transcript—but not SLOW-1 protein—is necessary to guarantee the correct expression of *slow-1* in the germline of its progeny.

### Licensing of *slow-1* is not equivalent to RNAa

We wondered whether mechanisms previously proposed to guarantee expression of endogenous genes—PATCs and CSR-1—might underlie epigenetic licensing of endogenous genes by maternal transcripts. We found that both *slow-1* and slow-2 had a very low PATC score in their intronic sequences, suggesting that PATCs are not involved in epigenetic licensing (Figure S12). CSR-1 binds chromatin in a 22G-RNA-dependent manner [28]. To understand the role of 22G-RNAs in licensing, we isolated and sequenced total sRNAs from both NIC203 and EG6180 parental strains. The total amount of 22G-RNAs derived from each toxin was drastically different. 22G-RNAs derived from *slow-1* were 10 times more abundant than those derived from *slow-2* (Figure 4A; p=0.042). As a control, no global differences in 22G-RNA abundance were observed between NIC203 and EG6180 (Figure 4A). In addition, 22G-RNAs were heavily biased towards the 5’ of *slow-1*; 61% of *slow-1* 22G-RNAs were derived exclusively from its first exon (Figure 4A). Given that these sRNAs were isolated from parental strains that express either *slow-1* or *slow-2*, our data suggests that *slow-1-*derived 22G-RNAs are mechanistically related to its epigenetic licensing.

**Figure 4.**
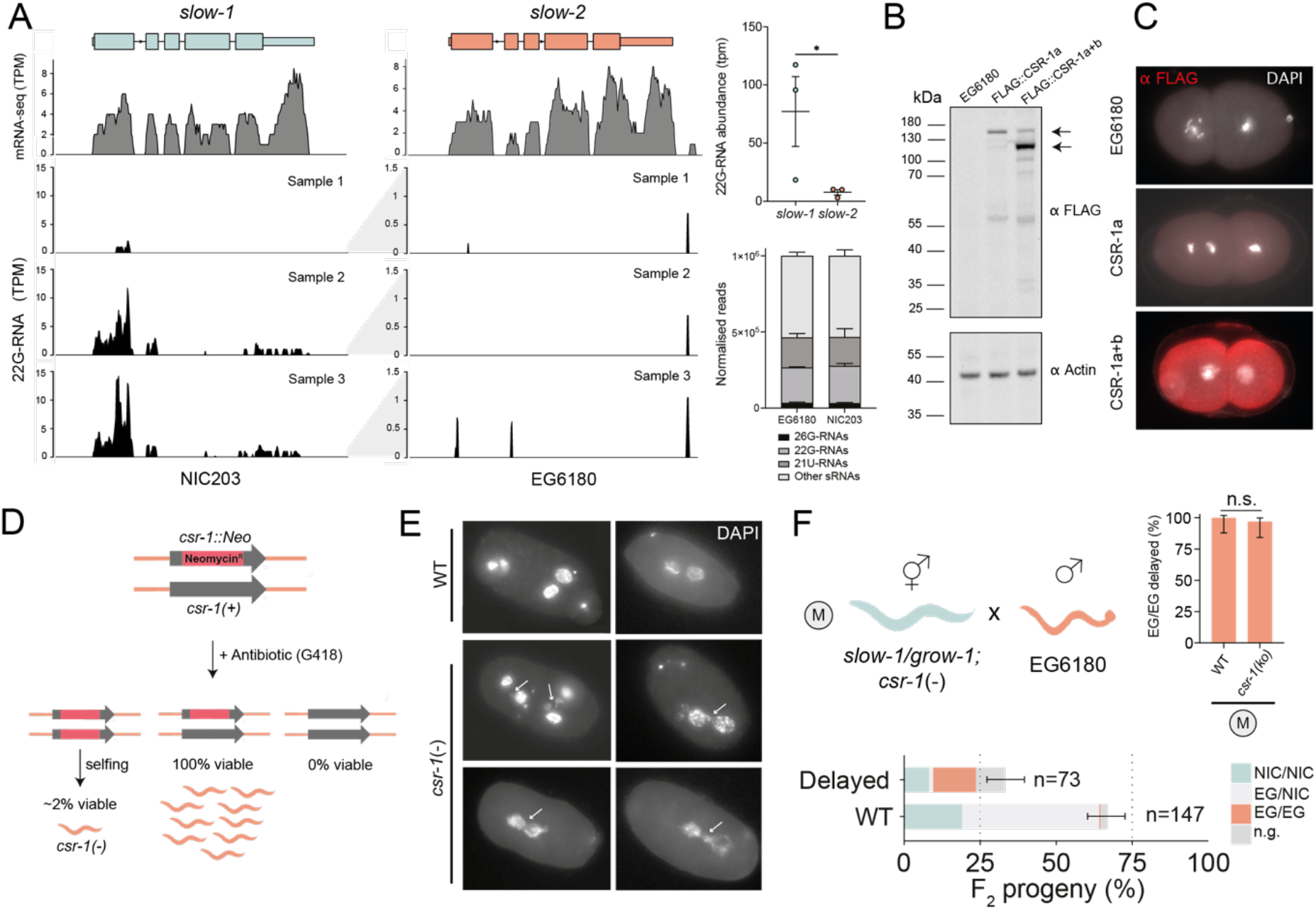
*slow-1* licensing has a 22G-RNA signature but is not equivalent to “RNAa”. **(A)** RNA-seq and 22G-sRNA short-reads aligned against *slow-1* and *slow-2*. sRNA libraries were generated in biological triplicates (left). Notice the differences in the y-axis scale between *slow-1* and *slow-2*. Quantification of total 22G-RNAs derived from *slow-1* and *slow-2* (top right, t-test p=0.042). Distribution of sRNA per parental strain (bottom right). **(B)** Detection of endogenously tagged FLAG::CSR-1a and FLAG::CSR-1a+b by western blot. Black arrows indicate the expected MW of each isoform. Western blot against β-Actin serves as a loading control. **(C)** Representative immunostaining images against FLAG::CSR-1a and FLAG::CSR-1a+b line in 2-cell stage embryos. EG6180 serves as a negative control. **(D)** Generation of balanced *csr-1* null strain by inserting *NeoR* in *csr-1* and growing worms in G418 antibiotic. **(E)** Representative images of wild-type (top), and *csr-1(-)* (bottom) embryos. Anaphase bridging events were observed in 14.75% of *csr-1(-)* early embryos (n=61) compared to 0% (n=36) of EG6180 WT. (**F)** Crosses between *csr-1*(-); *slow-1/grow-1* hermaphrodites and EG6180 males indicate that slow-1/grow-1 is fully active when maternally inherited. *csr-1(-)* hermaphrodites were derived from selfing of *csr-1(+/-)* mothers.

Given the 22G-RNA signature, we wondered whether epigenetic licensing depends on CSR-1. *C. elegans* codes for two CSR-1 isoforms: a long one, CSR-1a, which is expressed in spermatogenic gonad and a shorter one, CSR-1b, which is constitutively expressed in the female gonad [29, 30]. We identified a single homolog of *csr-1* in the genome of *C. tropicalis.* Tagging of the endogenous *csr-1* with either an N-terminal or an internal FLAG epitope revealed that both long and short isoforms are also expressed in *C. tropicalis* and immunofluorescence of early embryos was consistent with their known expression pattern in *C. elegans* (Figure 4B and C and Figure S13). To test whether CSR-1 is required for *slow-1* epigenetic licensing, we engineered a *C. tropicalis csr-1* null mutant disrupting both isoforms in the EG6180 background. Because *csr-1* is essential for viability in *C. elegans*, we first devised a strategy to stably propagate a *csr-1* heterozygous line in the absence of classical genetic balancers. To do so, we used CRISPR/Cas9 to introduce a premature stop mutation in the endogenous *csr-1* locus followed by a *neoR* cassette, which confers resistance to the G418 antibiotic (Figure 4D). We then propagated the mutant line by growing the worms in plates containing G418 and thus actively selecting for heterozygous *csr-1(-)* null individuals. Upon drug removal and genotyping, we found that most homozygous *csr-1(-)* individuals derived from heterozygous mothers developed into adulthood but were either sterile or laid mostly dead embryos. However, a small fraction of null mutants was partially fertile and homozygous csr-1(-) lines could be stably propagated for multiple generations despite extensive embryonic lethality in the population (Figure 4D). In agreement with the role of CSR-1 in *C. elegans* [28], DAPI staining of *csr-1(-)* null embryos revealed chromosome segregation problems, such as anaphase bridging events (Figure 4E), suggesting that CSR-1 plays a similar role in both species. In *C. elegans*, ‘RNAa’ is abolished in the F_1_ progeny if either of the parents is a heterozygous carrier of a null *csr-1* allele [15]. Thus, to test whether maternal *csr-1* is necessary for licensing in *C. tropicalis*, we crossed *csr-1*(-); *slow-1/grow-1* hermaphrodites to EG6180 males (see methods) and inspected their F_2_ progeny (Figure 4F). We found that 96.8% (n=32) of EG/EG F_2_ individuals were developmentally delayed, thus indicating that epigenetic licensing was not impaired in the absence of maternal CSR-1. Unfortunately, although a small fraction of *C. tropicalis csr-1(-)* hermaphrodites was viable and fertile, the high incidence of abnormal phenotypes in these worms precluded us from confidently testing the combined effect of maternal and zygotic *csr-1* loss of function on *slow-1* licensing. Overall, our results indicate that even though ‘RNAa’ and epigenetic licensing share similarities, they are not equivalent phenomena with regards to their dependency on CSR-1.

### *slow-1* is transgenerationally repressed

In *C. elegans*, silencing of transgenes can result in the stable inheritance of the repressed state for multiple generations. Typically, this transgenerational effect is mediated by heritable sRNAs in response to external (RNAi) or internal cues (piRNAs). To test whether sRNAs could underlie the impaired expression of the paternally inherited *slow-1/grow-1* allele, we asked whether the inheritance of *slow-1/grow-1* through the paternal lineage could compromise its toxicity in subsequent generations. To do this, we first crossed EG6180 hermaphrodites to NIL males and singled their F_2_ progeny. Then, we identified F_2_ homozygous *slow-1/grow-1* hermaphrodites, let them self-fertilize, collected their progeny (F_3_), and crossed them back to EG6180 males. Lastly, we collected the F_4_ heterozygous offspring, let them self-fertilize, and inspected their progeny (F_5_) (Figure 5A). In this way, the impaired *slow-1/grow-1* allele was reintroduced into the maternal lineage, which allowed us to probe whether *slow-1* could delay its progeny once again. We found that 97% (n=34) of F_5_ homozygous *slow-1/grow-1* (EG/EG) individuals were phenotypically wild-type, indicating that *slow-1/grow-1* activity was impaired three generations following paternal inheritance (Figure 5B). We then tested whether *slow-1/grow-1* was still inactive nine generations following paternal inheritance (Figure 5B). We observed that 22.2% (n=27) of homozygous EG/EG individuals were phenotypically wild-type, indicating that the repressed state could be reversed but was overall epigenetically stable (Figure 5B). In summary, when inherited through the paternal lineage and in the absence of maternal licensing, the *slow-1/grow-1* inactive state is transgenerationally inherited for multiple generations. Importantly, re-introduction of the allele into the maternal lineage is not sufficient for immediate reactivation and like in most cases of transgenerational inheritance in nematodes, the repressive state can be spontaneously reverted [31, 32].

**Figure 5.**
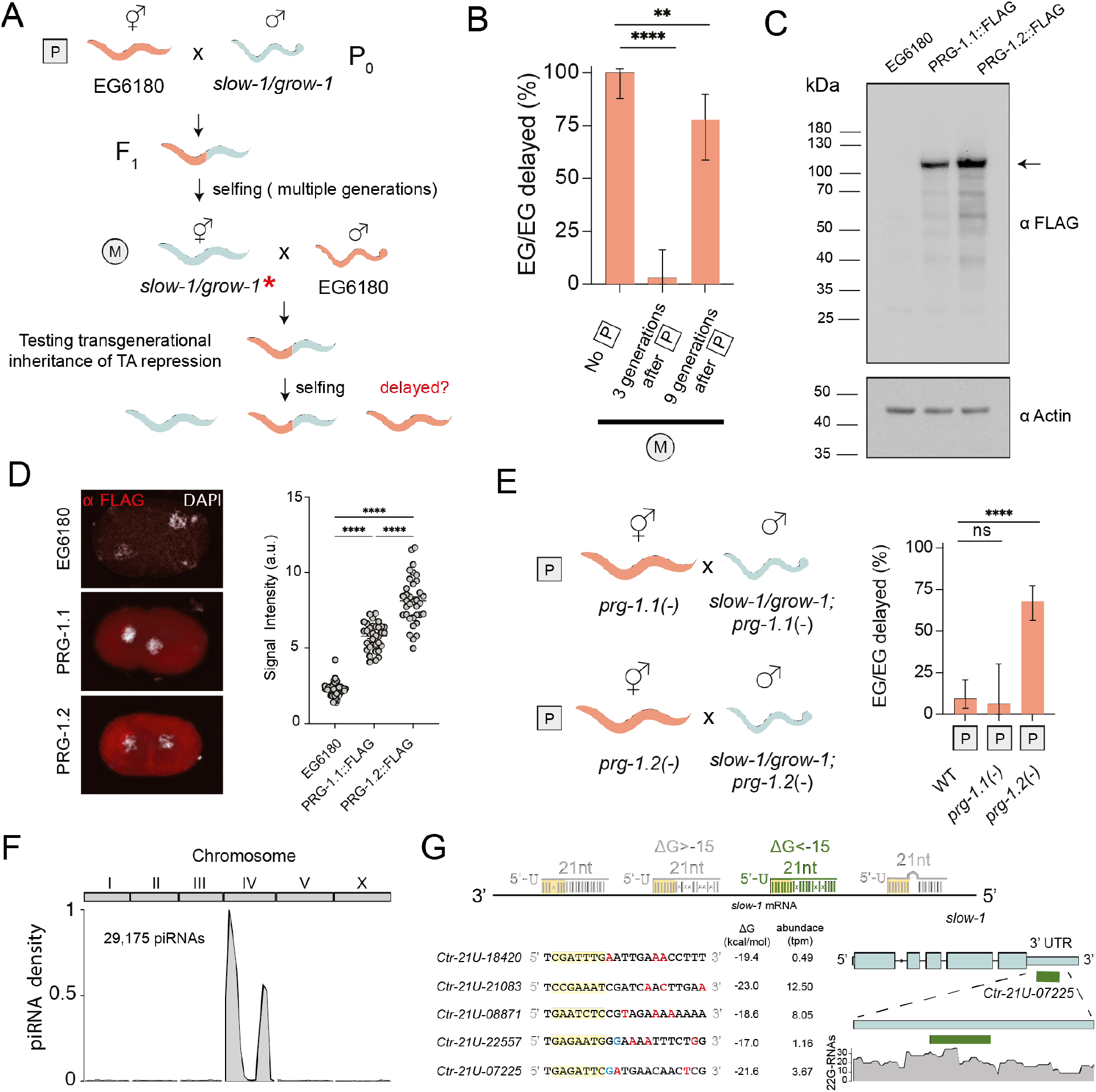
*slow-1* is transgenerationally silenced and targeted by the piRNA pathway. **(A)** Crossing scheme to test whether *slow-1/grow-1* repression is transgenerationally inherited. Initially, EG6180 hermaphrodites were crossed to NIL males, leading to silencing of *slow-1/grow-1* in the F_1_ progeny. F_2_ homozygous TA carriers were identified and propagated via selfing for three or nine generations and then crossed back to EG6180 males. The F_2_ progeny from these parents were then phenotyped for delay. **(B)** Comparison of *slow-1/grow-1* activity in the absence of paternal inheritance, three generations following paternal inheritance, and nine generations following paternal inheritance (n_ctl_=34, n_3gen_=34, n_9gen_=27, Fisher’s exact test comparison to control, p<0.0001 and p=0.0053). **(C)** Detection of endogenously tagged PRG-1.1::FLAG and PRG-1.2::FLAG by western blot. Black arrow indicates the expected MW. EG6180 was used as a negative control. Western blot against β-Actin serves as a loading control. **(D)** Representative immunostaining images against PRG-1.1::FLAG and PRG-1.2::FLAG line in 2-cell stage embryos (left). EG6180 serves as a negative control. Quantification of immunostaining (rigth) **(E)** Effect of either *prg-1.1* or *prg-1.2* null mutations in *slow-1/grow-1* paternal inheritance. Crosses between *prg-1.1*(-) hermaphrodites and NIL *prg-1.1*(-) males result in mostly wild-type EG/EG F_2_ individuals (n_ctl_=53, n_prg-1.1(-)_=16, Fisher’s exact test p>0.99). However, crosses between *prg-1.2*(-) hermaphrodites and NIL *prg-1.2*(-) males result in delayed EG/EG F_2_ individuals, indicating that *prg-1.2* is involved in the silencing of *slow-1* (n_ctl_=53, n_prg-1.2(-)_=75, Fisher’s exact test p<0.0001). **(F)** Genome-wide distribution of *C. tropicalis* piRNAs (21U-RNAs). Two large clusters are found in Chr. IV. **(G)** Computational identification of candidate piRNAs that bind *slow-1* mRNA (see Methods). Highlighted in yellow is the seed region. Mismatches between piRNAs and *slow-1* in red. Wobble base pair mismatches in blue.

### The piRNA pathway targets *slow-1*

Since the transgenerational repression of *slow-1/grow-1* does not stem from an external trigger, we reasoned that endogenous piRNAs could mediate this transgenerational effect. PRG-1, the *C. elegans* ortholog of *Drosophila* PIWI-clade proteins, is essential for piRNA function [8, 33]. The *C. tropicalis* genome codes for two PRG-1 orthologs on Chr. I, which we named PRG-1.1 and PRG-1.2. They share 87.7% sequence identity at the protein level and are likely the result of a recent gene duplication event (Figure 5C, D and Figure S14A; not to be confused with PRG-2, a *C. elegans* pseudogene). To test whether *slow-1/grow-1* repression was dependent on piRNA activity, we first generated *prg-1.1* and *prg-1.2* null alleles in an EG6180 background using CRISPR/Cas9. Both *prg-1.1* and *prg-1.2* mutant lines were viable and did not show any obvious signs of developmental delay or larval arrest (0%, n=118 and 0%, n=95 respectively). However, *prg-1.1 prg-1.2* double mutants were sterile indicating that these two genes are redundant and essential in *C. tropicalis* (Figure S14B). To test whether piRNAs are downregulated in these mutants, we first annotated piRNAs from EG6180 sRNA libraries and then quantified their abundance in both mutants. Overall, we identified 29,175 piRNAs in *C. tropicalis* with a mean abundance of 0.1 ppm or higher (Excel Table2). Like in *C. elegans* and *C. briggsae*, piRNAs almost exclusively derived from Chr. IV (96.9%; Figure 5F) [10]. Quantification of sRNA libraries using microRNA abundance for normalization, revealed that piRNAs were globally downregulated in *prg-1.1* and *prg.1.2* mutants, albeit partially (mean fold change 0.56 and 0.50, respectively), as expected from their redundant role.

Next, we set up crosses between EG6180 hermaphrodites and NIL males, in which both parents carried null alleles of either *prg-1.1* or *prg-1.2*. Strikingly, loss of *prg-1.2* impaired *slow-1/grow-1* repression when inherited through the paternal-lineage, 69.2% (n=78) of F_2_ homozygous EG/EG individuals were developmentally delayed, indicative of an active paternally inherited SLOW-1 toxin (Figure 5E). In contrast, loss of *prg-1.1* had only a minor effect on the activity of the TA (6.25% of EG/EG were delayed, n=16) (Figure 5E). We then tested whether *prg-1.2* was required maternally or zygotically to repress *slow-1*. When *prg-1.2* null mutant mothers were crossed to wild-type NIL males, we observed that 52.1% (n=23) of F_2_ homozygous EG/EG individuals were delayed. In contrast, when EG6180 mothers were crossed to *prg-1.2* null mutant NIL males, F_2_ homozygous EG/EG individuals were mostly wild-type (13.8% delay; n=29) (Figure S14C and D). Thus, maternal activity of *prg-1.2* is required to repress the paternally inherited *slow-1/grow-1* allele in the germline of the F_1_. Lastly, we evaluated whether *C. tropicalis* piRNAs could bind *slow-1* by leveraging known targeting rules and predicted piRNA-target binding energies (see methods). We identified five candidate piRNAs that could target *slow-1* but not *slow-2* (Figure 5G). Among the top candidates was *Ctr-*21ur-07225. This piRNA is abundant (3.67 ppm, top 25%) and predicted to bind the 3’ UTR of *slow-1*—it matches perfectly within the seed region and it has only three mismatches outside it, one of them, a G:U wobble base pair (ΔG=-21.6 kcal/mol). Moreover, *Ctr-*21ur-07225 showed the distinctive pattern of 22G-RNA accumulation associated with PRG-1 activity (Figure 5G) [34]. In summary, these experiments show that when piRNAs are impaired, the licensing activity of *slow-1* is dispensable for the activation of *slow-1* in the zygotic germline. In other words, the positive effect of epigenetic licensing on gene expression stems from inhibiting the repressive action of piRNAs (Figure 6).

**Figure 6.**
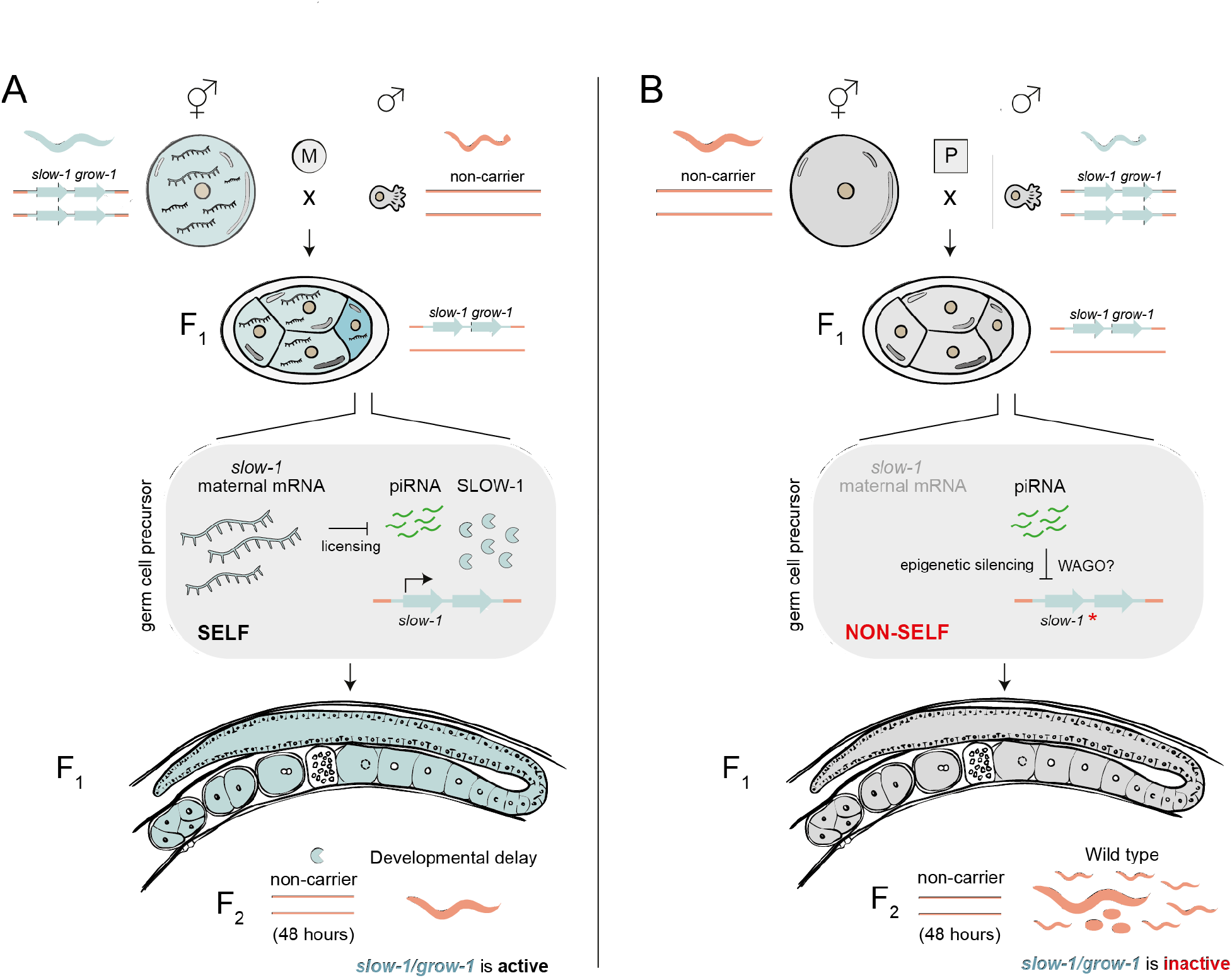
Model illustrating *slow-1/grow-1* parent-of-origin effect and epigenetic licensing. **(A)** Maternal-inheritance of the *slow-1/grow-1* TA. *slow-1* transcripts deposited in the egg by the mother are sufficient and necessary to activate zygotic *slow-1* in the germline of the F_1_ progeny. Epigenetic licensing stems from inhibiting the repressive action of piRNAs. F_1_ heterozygous mothers load SLOW-1 toxin into all of their eggs. F_2_ homozygous non-carrier individuals are developmentally delayed because they do not express the zygotic antidote. **(B)** Paternal-inheritance of the *slow-1/grow-1* TA. In the absence of *slow-1* maternal transcripts, piRNAs repress the transcription of *slow-1* in the germline of heterozygous F_1_ mothers. As a result, SLOW-1 levels are insufficient to poison F_2_ homozygous non-carrier progeny. The repressed state of *slow-1(*)* is transgenerationally inherited for over five generations.

## Discussion

We discovered a translation-independent role of maternal transcripts in counteracting the spread of selfish genes. The epigenetic licensing we present here has only been described for one gene to date: the *C. elegans* sex-determining gene *fem-1* (See Supplementary Discussion) [35]. In the absence of maternal *fem-1* transcripts, the paternal *fem-1* allele is not correctly activated by the zygote, which in turn results in reduced spermatogenesis and feminization of their gonads [35]. The authors hypothesized that this mechanism could establish a transgenerational memory of self, such that genes with no prior history of germline expression—viruses and transposons— would be specifically silenced while endogenous genes would be protected. However, in over a decade since this discovery, no other gene has been shown to be epigenetically licensed and it was not clear which selective forces might be responsible for the evolution of such a mechanism. While studying how selfish TAs spread in natural populations of *C. tropicalis*, we discovered that a selfish toxin, *slow-1*, is epigenetically licensed and, because of this, the *slow-1/grow-1* TA is inactive when paternally inherited. Thus, our results indicate that epigenetic licensing is evolutionary advantageous as it protects the host from selfish genes that rely on mating to spread in populations. Given that *fem-1* and *slow-1* share this unique mechanism of gene regulation despite their disparate molecular roles in two different species, we propose that epigenetic licensing is a common feature among germline-expressed genes. Licensing has probably gone largely unnoticed because mutations that remove all traces of mRNA are seldom studied.

We also found that a second toxin, *slow-2* is not epigenetically licensed. *Slow-1* and *slow-2* quickly diverged from an endogenous nuclear hormone receptor gene following a gene duplication event [20]. It is unclear at this point whether epigenetic licensing was a feature of their common ancestor that was lost in *slow-2* or whether it evolved exclusively in *slow-1* in response to its parasitic nature. However, given that *fem-1* is not a selfish gene, we favor the former scenario. In other words, we propose that *slow-1* and *slow-2* are at different stages in their transition from friend to foe—*slow-1* still has the epigenetic licensing of the ancestral gene, while *slow-2* has lost epigenetic licensing, giving it an advantage by allowing it to spread via males.

Future studies may further resolve how precisely maternal transcripts counteract piRNAs. In animals and plants, noncoding RNAs can exert a regulatory role by competing for miRNA binding sites, so called “molecular sponges” [36]. Our data, however, does not support *slow-1* acting as a piRNA sponge (Figure S9). Instead, licensing likely involves a catalytic step in which the maternal mRNA is processed and used as a template for the generation of sRNAs (Figure 4A). CSR-1 has been put forward as a key factor opposing the silencing activity of piRNAs [15,16,28,37–39]. We found that maternal CSR-1 is completely dispensable for *slow-1* licensing (Figure 5F); however, we cannot exclude the possibility that CSR-1 acts in a parallel pathway or that it plays a different role in *C. tropicalis*. Our work also provides insights into piRNA function. In transgenic lines, the effect of piRNAs on gene expression seems binary: reporters are either expressed or silenced. In sharp contrast, we found that piRNAs can fine tune the expression of endogenous genes in agreement with recent reports [34,40,41] and remarkably, this quantitative effect on gene expression can be transgenerationally inherited.

Epigenetic licensing provides a physiological framework to better understand intriguing phenomena such as mating-induced silencing and target-dependent siRNA suppression [26, 42]. It also shares similarities with transcriptional adaptation, a mechanism by which mutations that trigger mRNA degradation cause the upregulation of paralogous sequences in *C. elegans*, zebrafish and mice [43–46]. Both rely on homology to activate their targets and involve sRNA biogenesis. It is tempting to speculate that they could be related mechanisms acting in the germline and soma, respectively. Lastly, our work illustrates how a parent-of-origin effect can stem from the repressive action of sRNAs. Interestingly, genomic imprinting of the *Rasgrf1* locus in mice also depends on the activity of piRNAs, which target a retrotransposon linked to its differentially methylated region [47]. Our findings suggest that conflict between the host and selfish genes may be a fundamental driving force in the evolution of parent-of-origin effects, irrespective of whether the imprinted loci are the subject of sexual or parent-offspring conflicts.

## Acknowledgments

Research in the Burga lab is supported by the Austrian Academy of Sciences, the city of Vienna, and a European Research Council (ERC) Starting Grant under the European 20 Union’s Horizon 2020 program (ERC-2019-StG-851470). A.K. is supported by a Boehringer Ingelheim Fonds (BIF) PhD Fellowship. S.W. is supported by the European Union’s Framework Programme for Research and Innovation Horizon 2020 (2014-2020) under the Marie Curie Skłodowska Grant Agreement Nr. 847548. Work by the Ben-David lab for this study was supported by the Israel Science Foundation (grant no. 2023/20). We thank Life Science Editors for editing services. Sequencing data are available under NCBI project PRJNA850171.

## Material and Methods

### Maintenance of worm strains

Nematodes were grown on modified nematode growth medium (NGM) plates. The modified NGM contains 1% agar/0.7% agarose as *C.tropicalis* can burrow into the normal NGM. All the experiments involving the *C. tropicalis slow-1/grow-1* and *slow-2/grow-2* TAs were performed at 25°C. The experiments involving the NIC203 ChrII TA and *C. elegans sup-35/pha-1* toxin were performed at 20°C. *csr-1*(+/-) strains were grown in 6 cm NGM plates supplemented with 500 μl of G418 (25mg/ml) to select for heterozygous null individuals. All strains used in this study are available on Table S1. Some of the strains were provided by the CGC, which is funded by the NIH Office of Research Infrastructure Programs (P40 OD010440).

### Generation of *C. tropicalis* transgenic lines

For CRISPR/Cas gene editing, we adapted previously described protocols [48]. In brief, 250ng/µl Cas9 or Cas12a (IDT) proteins were incubated with 200ng/µl crRNA (IDT) and 333ng/µl tracrRNA (IDT) at 37°C for 10 minutes before adding 2.5ng/µl co-injection marker plasmid (pCFJ90-mScarlet-I). For HDR, donor oligos (IDT) or biotinylated and melted PCR products were added at a final concentration of 200ng/µl or 100ng/µl, respectively. Following injections into young hermaphrodites, mScarlet-positive F1 were singled, and their offspring screened by PCR and Sanger sequencing to detect successful editing events. To clone the mScarlet::SLOW-1 donor, we added around 300bp homology arms amplified from QX2345 genomic DNA to mScarlet-I (from pMS050) in pBluescript via Gibson assembly. For the *csr-1::neoR* donor, we first replaced the *C.elegans rps-27* promoter and *unc-54* 3’UTR in pCFJ910 with 500bp upstream and 250bp downstream of the *C.tropicalis rps-20* gene. This *rps-20::neoR* cassette was then flanked with ∼550bp homology arms amplified from EG6180 worms and inserted into pBluescript. Correct targeting introduces a stop codon after residue L337 of CSR-1 followed by a ubiquitously expressed neomycin resistance. All gRNAs and HDR templates are available on Table S3 and S4.

### Phenotyping and genotyping of crosses

To set up crosses, 4-5 L4 hermaphrodites were mated with around 30-40 males in a 12-well plate containing modified NGM. After 2 days, approximately 10 L4 F_1_ progeny were transferred to separate plates and allowed to lay embryos. The next day, at least 10 embryos per F_1_ hermaphrodite were singled into 6 cm NGM plates. In the absence of genetic markers to distigh cross-progeny from self-progeny, F_1_ hermaphrodites were genotyped using PCR. F_2_ individuals from self-progeny were excluded from the final analysis. Each F_2_ individual was visually inspected once per day at approximately 24 hours intervals and scored for developmental stage and any phenotypic abnormality for up to 7 days. In cases where after 24 hours the embryo did not hatch, it was classified as embryonic lethal. If after 7 days the worm did not reach adulthood, it was classified as arrested. In cases where the F_2_ individual took 3 days or more to produce offspring, it was marked as delayed. For simplicity, in this manuscript we use the term “delayed” to include all individuals that are classified as delayed plus a small percentage of arrested individuals (0-20% of affected individuals). If the worm was an adult but did not produce progeny, it was classified as sterile. After 7 days, at least 10 worms per plate (whenever possible) were placed on a single PCR tube containing lysis buffer (50 mM KCl, 10 mM Tris pH 8.3, 2.5 mM MgCl2, 20mg/ml proteinase K), lysed (30 min at 60℃ followed by 15 min at 95℃) and genotyped by PCR followed by agarose gel or sanger sequencing. A list of primers used for genotyping can be found here (Table S2). Importantly, all phenotypic scores were done before knowing the genotype of the F_2_. For crosses between *csr-1*(-); *slow-1/grow-1* hermaphrodites vs EG6180 males, males carried a pmyo-2::mScarlet single copy reporter in the EG6180 background (INK351). For crosses between injected hermaphrodites and NIL males, males also carried the pmyo-2::mScarlet reporter (INK460). In both cases, only fluorescent F_1_ hermaphrodites were selected for further experiments.

### *In vitro* RNA transcription and injection

To prepare for in vitro RNA transcription, the *slow-1* cDNA was cloned into pGEM-T Easy (Promega, #A1360) backbone, with a 5’-T7 RNA polymerase site and the start codon mutated to prevent translation of the RNA (ATG>TTG). The plasmid was transformed and midi-prepped (Qiagen, #12143), then the insert was isolated by digesting approximately 10ug of plasmid with NotI (NEB, #R0189) for 2hrs at 37°C, followed by gel extraction. The RNA was transcribed using 1ug of template with the HiScribe T7 Quick High Yield kit (NEB, E2050) according to the manufacturer’s instructions with the following modifications: 3µL of 10mM DTT was added, as well as 1µL of RNaseOUT (Thermo, 10777019). The transcription reaction was allowed to run overnight to maximize RNA output. Then the reaction was diluted to 50µL and 2µL RNase-free DNase I (NEB, M0303S) were added. The DNase I digest was performed at 37°C for 15 minutes. The RNA was then bead-purified (Vienna Biocenter MBS 5001111, High Performance RNA Bead Isolation), quantified using the Qubit RNA HS Assay kit (Thermo, Q32852), run on a gel to check for RNA integrity and stored at −80°C. The injections were repeated twice using RNA independently transcribed and using two different concentrations: 150nM and 400nM with identical results.

### Reciprocal crosses with the *mScarlet::slow-1* reporter line

To measure the expression of SLOW-1 in the F_1_ progeny of the reciprocal crosses between mScarlet::SLOW-1 NIL and EG6180 strains, we set up the following crosses: 1) SLOW-1::mScarlet dpy (INK461) hermaphrodites to EG6180 males to study the SLOW-1 expression when inherited maternally and 2) EG6180 *dpy* (QX2355) hermaphrodites to mScarlet::SLOW-1 NIL males (INK459) when paternally inherited. We first immobilized wild-type young adult F_1_ progeny in a drop of NemaGel on a glass slide and coverslipped it. Immediately after immobilization, images were acquired using an Axio Imager.Z2 (Carl Zeiss) widefield microscope equipped with a Hamamatsu Orca Flash 4 camera, with an Ex 545/30nm filter. The analysis was performed in FIJI, by tracing the germline of each worm first in the DIC channel and later measuring the mean fluorescence. The fluorescence signal also includes gut autofluorescence.

### Sequencing and genome assembly of EG6180

We extracted high molecular weight (HMW) genomic DNA using the Masterpure Complete DNA and RNA purification kit (Lucigen), using the protocol for tissue samples, with the modification that worms were frozen and thawed to facilitate lysis prior to extraction. We then prepared 8 kb, 20 kb and unfragmented sequencing libraries using the 1D Ligation Sequencing Kit from Oxford Nanopore Technologies (SQK-LSK109). 8 kb was done using g-TUBE (Covaris) following manufacturer protocols. Library was loaded on a MinION MK1B device (Oxford Nanopore). Read calling was done using MinKNOW software. We carried out a hybrid assembly strategy as described in a previous study that also includes the Illumina sequencing reads of EG6180 used in this assembly with some differences (see below) [20]. N50 of Nanopore reads was 5,206bp. We used assembled Illumina reads to correct raw Nanopore reads, which were taken for assembly using Flye Assembler [49]. The preliminary assembly included 119 contigs in 107 scaffolds (Scaffold N50 was 1,489,504bp). We derived synteny blocks between the provisional assembly and our chromosome-level NIC203 assembly using Sibelia [50] and used the synteny blocks to scaffold the contigs to chromosome level using Ragout [51].

### Transgenerational silencing of *slow-1/grow-1*

For the transgenerational inheritance experiments, we first crossed EG6180 hermaphrodites to NIL (QX2345) males. Then, F_1_ individuals were isolated and genotyped after they had laid embryos to distinguish between self-progeny and cross-progeny. Then we singled F_2_ embryos from cross-progeny mothers, waited until they laid eggs and genotyped them. F_3_ homozygous carriers for slow-1/grow-1 were singled and propagated for multiple generations and mated to EG6180 males. We then estimated the activity of the *slow-1/grow-1* TA as previously described by determining the proportion of delayed EG/EG non-carriers.

### Single molecule *in situ* hybridization

Custom Stellaris FISH Probes were designed against *slow-1, slow-2 and pgl-1* using the Stellaris RNA FISH Probe Designer (Biosearch Technologies, Inc., Petaluma, CA) available online at www.biosearchtech.com/stellarisdesigner. The samples were hybridized with the Stellaris RNA FISH Probe set labeled with Quasar® 570, CAL Fluor Red 610 or Quasar® 670, respectively (Biosearch Technologies, Inc.), following the manufacturer’s instructions available online at www.biosearchtech.com/stellarisprotocols. The protocol was adapted from Raj et al [52] and described in [20]. For imaging, an Axio Imager.Z2 (Carl Zeiss) widefield microscope equipped with a Hamamatsu Orca Flash 4 camera was used. Images were acquired with a 63x/1.4 plan-apochromat Oil DIC objective with the following filters: for DAPI Ex 406/15nm, Em 457/50nm and for Quasar® 570 Ex 545/30nm, Em 610/75nm. For each image a Z-stack containing 40 slices with a step size of 0.2 um was acquired. For image analysis the FIJI plugin RS-FISH[53] was used with the following parameters: Sigma 1.44, threshold 0.0062.

### RNA extraction and mRNA-seq

We extracted total RNA from approximately 100 young adult hermaphrodites. To isolate RNA from F_1_ progeny, we made use of recessive mutations that allowed us to visually discriminate cross-progeny from self-progeny. Reciprocal crosses between parental strains were set up in the following way. To study maternal inheritance of *slow-1/grow-1*, we mated INK531 hermaphrodites (*uncoordinated* worms in the NIC203 background) to EG6180 males. To study paternal inheritance of *slow-1/grow-1*, we mated QX2355 hermaphrodites (*dumpy* worms in the EG6180 background) to NIC203 males. In both cases we only selected phenotypically WT progeny for RNA extraction. Reciprocal crosses between the NIL and EG6180 strains were performed in an analogous fashion. To study maternal inheritance of *slow-1/grow-1*, we mated INK255 hermaphrodites (*dumpy* worms in the NIL background) to EG6180 males. To study paternal inheritance of *slow-1/grow-1*, we crossed QX2355 hermaphrodites (*dumpy* worms in the EG6180 background) to QX2345 NIL males. Total RNA was extracted following a modified version of the protocol in [54]. Shortly, worms were washed multiple times using M9, followed by incubation with TRizol and chloroform. Phase-separating columns were used to separate the TRIzol and the aqueous phase was mixed with an equal volume of isopropanol. Following centrifugation, the RNA pellet was washed multiple times and then resuspended in RNase-free water. Only RNA samples with RIN>8 were used for library preparation using the NEBNext Poly(A) kit. The samples were sequenced on NextSeq2000 P2 SR100 or NovaSeq S1 PE100 at the Vienna Biocenter NGS facility. To reduce reference bias in our NIC203/EG6180 allele specific expression analysis, raw reads were aligned to a concatenated NIC203+EG6180 genome/transcriptome assembly with STAR, using the RNA-seq pipeline available in bcbio-nextgen (https://github.com/bcbio/bcbio-nextgen). Transcript quantification was carried out in tximport as implemented in bcbio-nextgen. Transcript counts were then normalized using DEseq2 [55]. We used DEseq2 to fit a model for the normalized counts using the strain identity of the mother and sequencing batch (Nextseq vs NovaSeq libraries) as fixed effects and compared the model to a null model that included only batch using a likelihood-ratio test, as implemented in DEseq2.

### Small RNA library preparation and sequencing

To isolate RISC associated sRNAs, we used the TraPR protocol [56]. In brief, frozen worm pellets (2000 worms for each parental line) were supplemented with 350µL lysis buffer (20mM HEPES-KOH, pH 7.9, 10% (v/v) glycerol, 1.5 mM MgCl_2_, 0.2mM EDTA, 1mM DTT, 0.1% v/v Triton x100, conductivity adjusted at 8 mS cm-2 with KoAc at an indicative concentration of 0.1M). Samples were then mechanically disintegrated and subjected to 4 freeze-thaw cycles in liquid nitrogen. Resulting lysates were cleared by centrifugation (10 000 x g, 5 min, 4℃). The sRNA fraction was isolated by TraPR Small RNA Isolation Kit (135.24, LEXOGEN) following manufacturers instructions. Isolated sRNA was treated with RNA 5’ Pyrophosphohydrolase (RppH, M0356S, BioLabs), to ensure 5’ monophosphate-independent capturing of small RNAs as previously described [57]. Following the enzymatic reaction, Agencourt RNA Clean XP magnetic beads (BECKMAN COULTER) were used to purify the sRNA. Purified sRNA was ligated to a 32 nt 3’ adaptor including unique barcodes (sRBC, Table S5, IDT) with truncated T4 RNA ligase 2 (M0373L, NEB) overnight at 16°C. The resulting RNA product was run on 12% SequaGel -UreaGel (national diagnostics) and purified with ZR small-RNA PAGE Recovery Kit^TM^ (R1070, ZYMO RESEARCH) following provider’s instructions. The 37nt long 5’ adaptor was ligated to the small RNAs using T4 RNA ligase (M0204S, NEB) overnight at 16°C. Enzymatic reaction was cleaned up using RNA Clean and Concentrator 5 kit (R1015, ZYMO RESEARCH). Ligated RNA was reverse-transcribed and PCR amplified. Resulting cDNA was separated on a 4% agarose gel and the 160–190 nt long fragments were extracted and gel purified using the Zymoclean Gel DNA Recovery Kit (D4008, ZYMO RESEARCH). Small RNA Libraries were sequenced in triplicates on a NovaSeq S1 SR100 mode (Illumina) at the Vienna Biocenter NGS facility.

### Small RNA analysis

5’ and 3’ adapters were trimmed using Cutadapt v1.18 [58]. 21U and 22G reads were extracted using a custom script and aligned to the genome using hisat2 v2.1 [59]. For 22G, only reads mapped to the coding sequences were analyzed; for 21U, reads mapped to coding sequences, tRNAs and rRNAs were excluded using seqkit v0.13 and samtools v1.10. 22G reads were quantified using featureCounts function of Rsubread R package, normalized by the total number of 22G per replicate, and visualized using the Gviz R package [60]. Of 21U reads, only ones that mapped perfectly and had an abundance more than 0.1 ppm (∼20-30 raw reads observed) were considered candidate piRNAs and used for further analysis. Quantification was performed by custom script, reads were normalized by miRNAs predicted based on homology to known *C. elegans* miRNAs. Nucleotide blast v2.2.26 in “blastn-short” mode against *slow* transcript was used to identify candidate piRNAs that could potentially bind *slow-1*. Next, binding sites and energies of those piRNAs were predicted with RNAduplex from the ViennaRNA Package v2.0 [61]. Candidate piRNAs that do not form bubbles during binding, with ΔG < −15, with 0 mismatches in the seed region, no more than 3 mismatches overall, and no more than 1 G·U wobble, were extracted by a custom script.

### Immunohistochemistry

Gravid nematodes cultured on three 9cm NGMA plates were recovered in 15 ml tubes by washing with M9 buffer. The nematodes were collected by centrifugation (500 rcf, 1min, RT) and once more washed with M9. To extract embryos from gravid adults, 500µl bleach solution (Ecolab Sator, 1plushygiene) and 500µl 1M NaOH were added to the worm pellets in 1.5ml of M9. The bleaching procedure was stopped immediately once dissolving of adults was observed by diluting in M9. Multiple washes with M9 followed. The embryo suspension was applied to a prepared poly-L-lysine slide (Sigma-Aldrich, P8920), shielded by a coverslip and immersed into liquid nitrogen for 15s. Subsequently, the coverslip was flicked off and the slides with embryos were fixed by an immediate incubation in ice cold methanol (15 min) followed by an incubation in ice cold acetone (10 min). The slides were then submitted to an ethanol rehydration row by a 5 min incubation in each of four descending ethanol concentrations (EtOH 95%, 70%, 50%, 30%). The fixation was finished by a single wash in 1x PBS (5 min). A nonspecific antibody binding was prevented by an incubation of the fixed embryos in 3% BSA (VWR Life Science, 422351S) in PBS-T (1h, RT). Binding of the anti-FLAG M2 primary antibody (mouse, Sigma-Aldrich, F3165, diluted 1:3000 in PBS-T+1%BSA) was allowed in a humidity chamber (overnight, 4°C). After 3 washes in PBS-T (7min each), a secondary antibody Alexa FluorTM A568 (Goat anti-Mouse, ThermoFisher Scientific, A-11031) was applied in dilution 1:3000 (1h, RT). Antibody stainings were followed by four washes in PBS-T (7 min each). Second PBS-T wash was enriched for DAPI (Merck, D9542, 5 ng/mL). Processed embryos were mounted with Fluoroshield (Sigma-Aldrich, F6182) and immediately imaged at Axio Imager 2 (ZEISS).

### Fluorescence intensity quantification

24-bit raw images were analyzed in Fiji (v1.53r) [62]. Embryos were selected by freehand tool and the same selection mask was used to capture background fluorescence intensity for each embryo. To compare fluorescence intensities between strains we used corrected total cell fluorescence parameter (CTCF=Integrated Density – (Area of selected cell x Mean fluorescence of background readings). At least 23 embryos were used for quantification.

### Worm protein lysate preparation and western blot

Gravid adult worms were washed off from three 9 cm NGMA plates with M9 solution and then washed twice in the same solution. Worm pellets were flash-frozen in the liquid nitrogen and stored at −70°C until protein extraction. Worm pellets were resuspended in 0,3 ml of ice-cold lysis buffer (30 mM HEPES pH7.4, 100 mM KCl, 2 mM MgCl2, 0.05% IGEPAL, 10% glycerol and 1 tablet of protease inhibitors (Roche, 11836153001) per 10 ml of solution and lysed by sonication in Bioruptor (UCD-200, Diagenode) with the following settings: H, 10 intervals with 30 seconds sonication, 30 seconds rest. Samples were incubated with rotation for 20 min at +4°C followed by two rounds of centrifugation (20000 rcf, 4°C, 10 min) to clear the supernatant. Protein concentration was measured by Bradford assay (Thermo Scientific, 23238). Samples were diluted to equal concentrations and resuspended in a 5X SDS loading buffer (0,25 M Tris pH6.8, 10% w/v SDS, 50% glycerol, 0,5 M DTT, 0,25% w/v Bromophenol Blue). 40 μg of each sample was loaded per well and proteins were separated by SDS polyacrylamide gel electrophoresis using precast NuPAGE gels (Invitrogen, NP0301BOX). Samples were transferred to 0,45 µm PVDF membrane (Thermo Scientific, 88518). The membrane was blocked with 4% non-fat milk in TBS-T (TBS, 0,1% Tween-20). After blocking, membranes were incubated with anti-FLAG M2 (mouse, 1:2000, Sigma-Aldrich, F3165) or anti-Actin (rabbit, 1:3000, Abcam, ab13772) primary antibody in a blocking solution overnight at 4°C. After primary antibody incubation membranes were washed three times with TBS-T for 5 min followed by incubation with HRP-conjugated anti-mouse (1:10000, Invitrogen, G-21040) or anti-rabbit (1:10000, Jackson Immuno, 111-035-045) secondary antibody in blocking solution for 1 hour at room temperature. Membranes were washed three times with PBS-T for 5 min followed by 1 minute incubation with ECL detection reagent (Cytiva, RPN2106) and imaged with ChemiDoc MP Imaging System (Bio-Rad). Membranes were washed twice in TBS and stripped before reprobing according to stripping reagent manufacturer’s protocol (Thermo Scientific, 21059).

### Live imaging of mScarlet::SLOW-1

For imaging, ∼20 gravid adults were dissected in M9 medium using a stereo microscope, and embryos were transferred to individual wells containing 70 µl of the same solution in a Thermo Scientific™ Nunc™ MicroWell™ 384-Well Optical-Bottom Plate (Thermo Scientific). Embryos were imaged using a Olympus spinning disk confocal based on an Olympus IX3 Series (IX83) inverted microscope, equipped with a dual-camera Yokogawa W1 spinning disk (SD) (Yokogawa Electric Corporation) and two ORCA-Flash4.0 V3 Digital CMOS cameras (Hamamatsu). Fields with suitable embryos were identified by performing a pre-scan of each well using a 10×/0.4NA (Air) objective. Each field was imaged by using a 40×/0,75NA (Air) objective and 16 *z*-sections at 2 µm. Imaging conditions were as follows: brightfield, 100% power, 30 ms; 568 nm, 100% power, 500 ms. All image acquisitions were performed using CellSense software (Olympus). Image processing and montages were created using Fiji and an automated cropping software, embryoCropUI [63].

## Supplementary Text

### The *slow-1/grow-1* TA has two redundant antidotes

We previously reported the existence of a maternal-effect TA in NIC203 Chr. III and identified the genes encoding the toxin and its cognate antidote, *slow-1* and *grow-1*, using CRISPR/Cas9-aided HDR repair [20]. While generating new mutant lines for the present study, we noticed that a *grow-1* mutant line was phenotypically WT despite carrying an active toxin. This result suggested the presence of an additional antidote. Upon careful inspection of the NIC203 *de novo* genome assembly and the underlying Nanopore raw reads, we discovered that there were two copies of the *grow-1* antidote instead of one. These two copies, which we named *grow-1.1* and *grow-1.2*, were identical at the nucleotide level and in close proximity, which likely caused an error in the genome assembler. In retrospect, we were fortunate while generating the original *grow-1* mutant in the *slow-1* null background. The *grow-1* gRNA we used targeted both copies and, unbeknownst to us, we had obtained a *grow-1.1* and *grow-1.2* double mutant. To facilitate our work with *grow-1.1* and *grow-1.2*, we designed gene specific primers pairs. Since these two genes are identical and redundant, we collectively refer to them as *grow-1* in this manuscript. In other words, all grow-1 mutants are *grow-1.1 grow-1.2* double mutants. On the other hand, *grow-2* only has one copy in the genome.

### Difference between a maternal-effect TA and parent-of-origin effects

TAs are made up of two linked genes: a toxin and its cognate antidote. Toxins are expressed in the germline of carriers, whereas antidotes are expressed zygotically. With the exception of the *C. elegans peel-1/zeel-1* TA, all other known TAs have a maternal-effect, that is, their toxic activity is transmitted through the maternal germline. This can be easily shown by performing reciprocal crosses between heterozygous TA carriers and homozygous non-carriers. In the case of maternal-effect TA, a cross between heterozygous mothers and homozygous males results in 50% affected progeny, whereas a cross between heterozygous males and homozygous mothers results in no affected progeny. It is also worth noting that in all maternal-effect toxins for which data is available, the toxins are loaded as protein into unfertilized eggs, analogously to maternal factors that are normally provisioned to ensure proper embryonic development. On the other hand, parent-of-origin effects occur when the phenotypic effects of a gene are determined by whether the gene is inherited from the mother or the father. In mammals, epigenetic imprinting is an example of a parent-of-origin effect. In contrast to all previously known TAs, the *slow-1/grow-1* TA has a parent-of-origin effect because it is only active when inherited through the maternal lineage. In summary, maternal and parent-of-origin effects describe two different phenomena. The first one is primarily concerned with the tissue where the toxin is expressed (either eggs or sperm), whereas the second one is concerned with differences in the activity or expression of the toxin between maternally and paternally inherited alleles.

### On epigenetic licensing and related terminology

In this manuscript, we used the term “epigenetic licensing” as originally coined by Johnson and Spence (2011) when studying *C. elegans fem-1*. The authors found that maternal transcripts of the sex-determining gene *fem-1* are required to license expression of a wild-type *fem-1* allele in the zygotic germ line. Thus, “epigenetic licensing” requires establishing a causal link between the presence of maternal transcript and the activation of the cognate zygotic gene. In later studies, the term “licensing” has also been used to describe the role of the Argonaute CSR-1 in counteracting piRNA silencing. For instance, Seth and colleagues (2013) found that RNAa “licenses” previously silenced transgenes and Wedeles and colleagues (2013) found that tethering of CSR-1 was sufficient to “license” their expression of silenced transgenes. However, it is important to mention that neither of these two studies established a link between this “licensing” activity and maternal transcripts. Importantly, whether CSR-1 activity is required for *fem-1* epigenetic licensing has not been shown to date. Moreover, a recent study that identified CSR-1 “targets” on a genome wide-scale failed to identify *fem-1* among them [38]. Thus, it cannot be assumed that “epigenetic licensing” by maternal transcripts is equivalent to “licensing” by CSR-1.

