## Supplement for "The evolution of an RNA-based memory of self in the face of genomic conflict"

**Supplementary Materials**

### Supplementary Figures

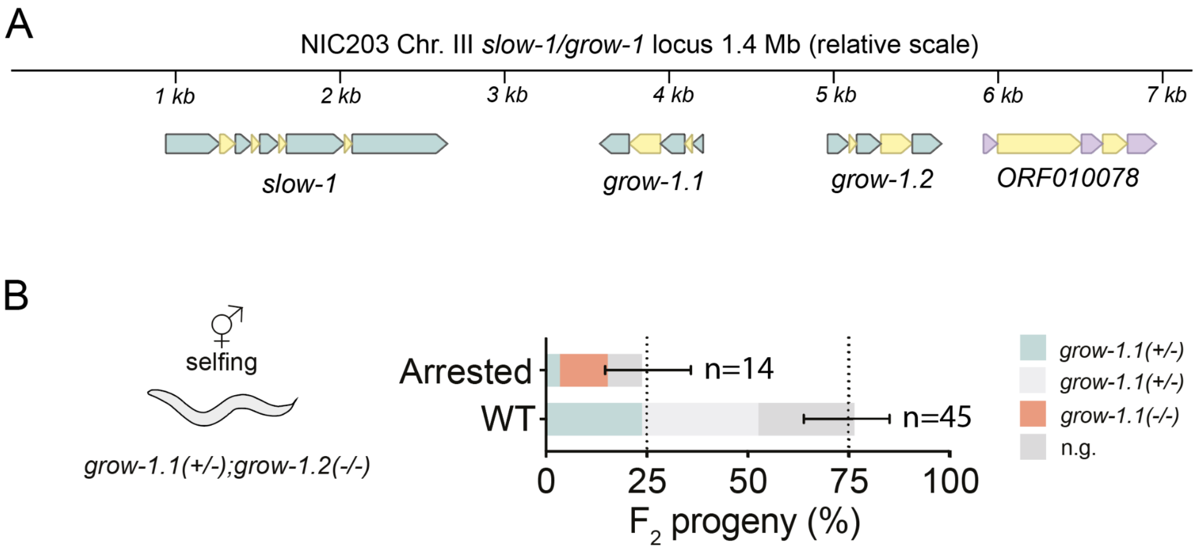


**Figure S1. The slow-1/grow-1 TA locus has two redundant antidotes. (A)** Corrected NIC203 genome assembly showing segmental duplication of the *grow-1* antidote **(B)** Selfing of *grow-1.1*(+/-); *grow-1.2*(-/-) strain. All *grow-1.1*(-/-); *grow-1.2*(-/-) individuals were developmentally arrested during larval development and did not produce any viable offspring. Thus, *grow-1.1* and *grow-1.2* are genetically redundant.

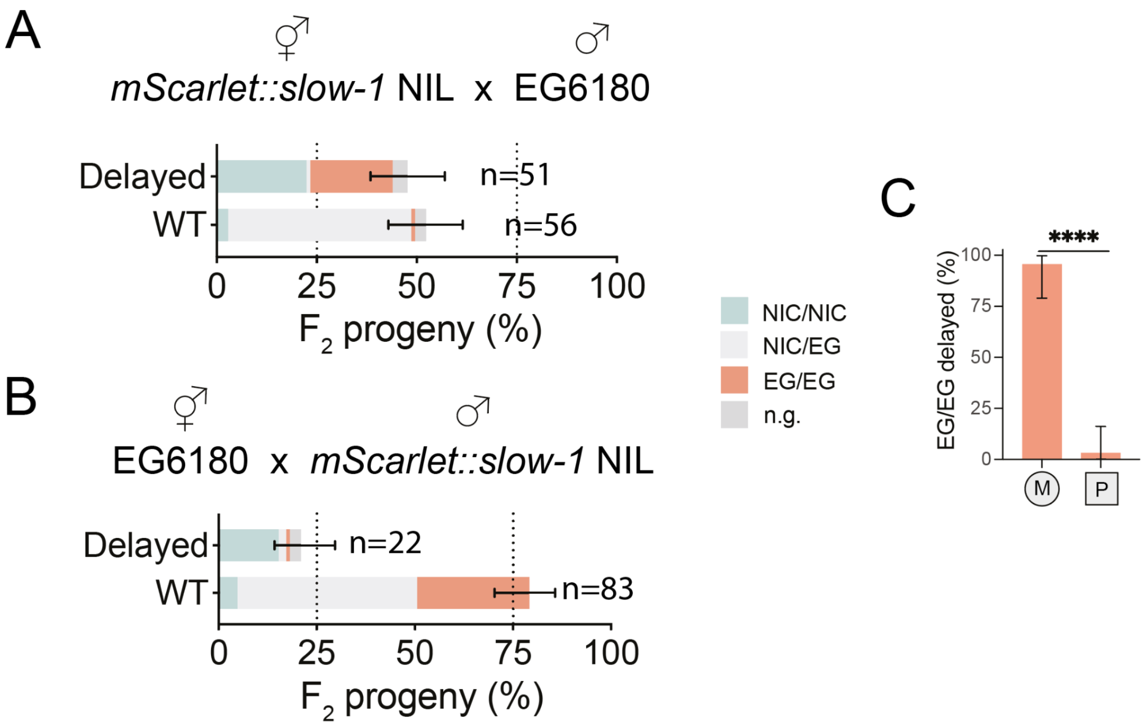


**Figure S2. The mScarlet::SLOW-1 endogenously tagged toxin is active and epigenetically licensed. (A)** A cross between mScarlet::SLOW-1 hermaphrodites and EG6180 males shows that the SLOW-1 fusion toxin is fully active, as indicated by the high percentage of $F_2$ EG/EG delayed progeny. **(B)** Cross between EG610 hermaphrodites and mScarlet::SLOW-1 males, shows that the SLOW-1 tagged toxin is not active when paternally-inherited, as observed for the *slow-1* WT allele. Notice that in both cases there are also $F_2$ NIC/NIC delayed progeny due to the presence of *slow-2/grow-2* locus in EG6180**. (C)** Percentage of $F_2$ EG/EG delayed progeny when *slow-1* is inherited maternally or paternally ($n_M$=23, $n_P$=31, Fisher's exact test $p<0.0001$).

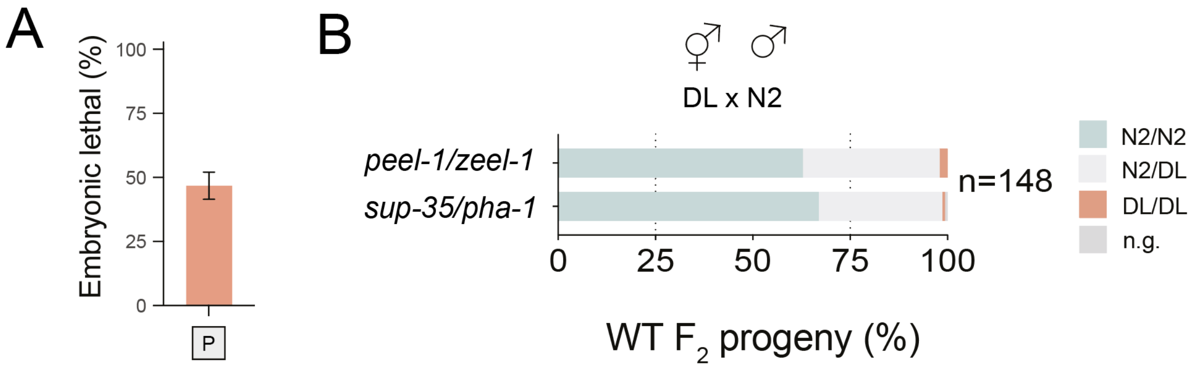


**Figure S3. The *C. elegans sup-35/pha*-1 TA is active when paternally inherited. (A)** Previously, we showed that *sup-35/pha-1* is active when maternally inherited (Ben-David, et al. Science 2017). To test whether *sup-35/pha-1* is active when paternally inherited, we crossed DL238 hermaphrodites and N2 males. N2 carries two TAs, *peel-1/zeel-1* and *sup-35/pha-1*. We observed 46.7% embryonic lethality among their $F_2$ progeny (n=340), as expected from the activity of two TAs segregating independently. **(B)** To confirm the activity of both TAs, we genotyped wild-type $F_2$ progeny for both *peel-1/zeel-1* (Chr. I) and *sup-35/pha-1* (Chr. III) and found that the vast majority of WT progeny were either homozygous or heterozygous carriers**,** indicating that both *peel-1/zeel-1* and *sup-35/pha-1* non-carrier individuals died as embryos.

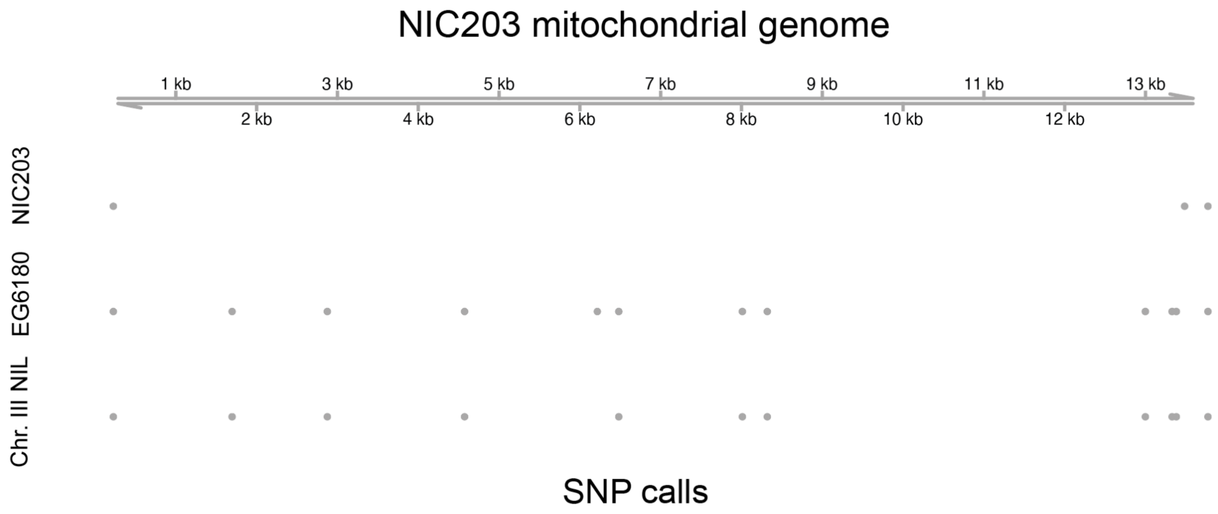


**Figure S4. The *slow-1/grow-1* Chr. III NIL strain and EG6180 have the same mitochondrial genotype.** Illumina short-reads from NIC203, EG6180, and Chr. III NIL DNA libraries were aligned against the NIC203 mitochondrial genome. Each dot represents a SNP. As expected from our cross scheme, the Chr. III NIL has the EG6180 mitochondrial genotype. Those SNPs shared by all strains likely reflect an error in the original NIC203 assembly.

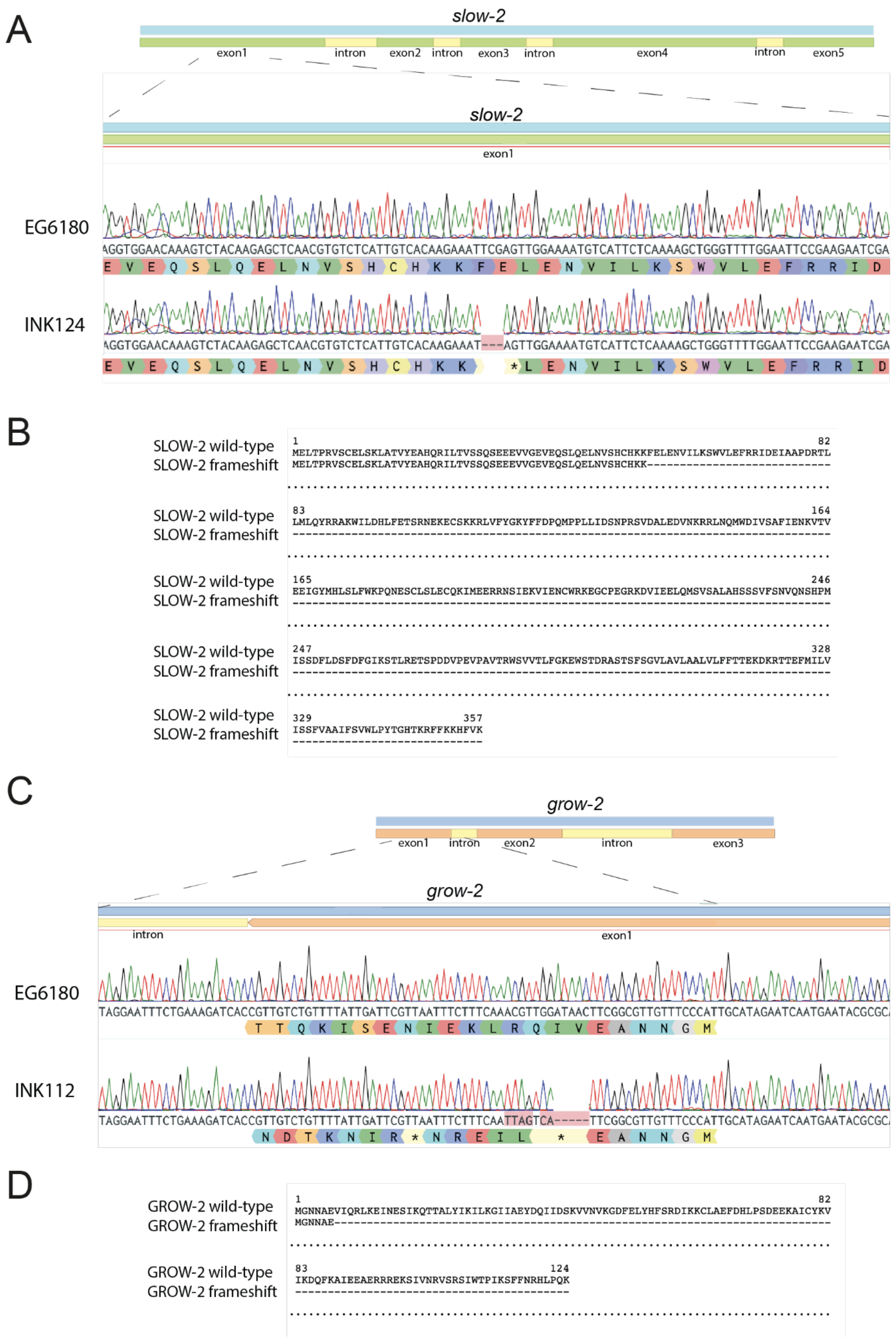


**Figure S5. Characterization of *slow-2* and *grow-2* null alleles. (A)** The *slow-2* null allele generates a three base pair deletion in the first exon, which creates a premature stop codon. This results in a shorter peptide (54 aa compared to 357 aa). **(B)** Protein alignment of wild-type SLOW-2 and mutant SLOW-2. **(C)** The *grow-2* null allele generates a deletion and frameshift in the first exon which creates a premature stop codon. This results in a shorter peptide (6 aa compared to 124 aa). **(D)** Protein alignment of wild-type GROW-2 and mutant GROW-2. Representative sanger sequences of WT and mutant alleles.

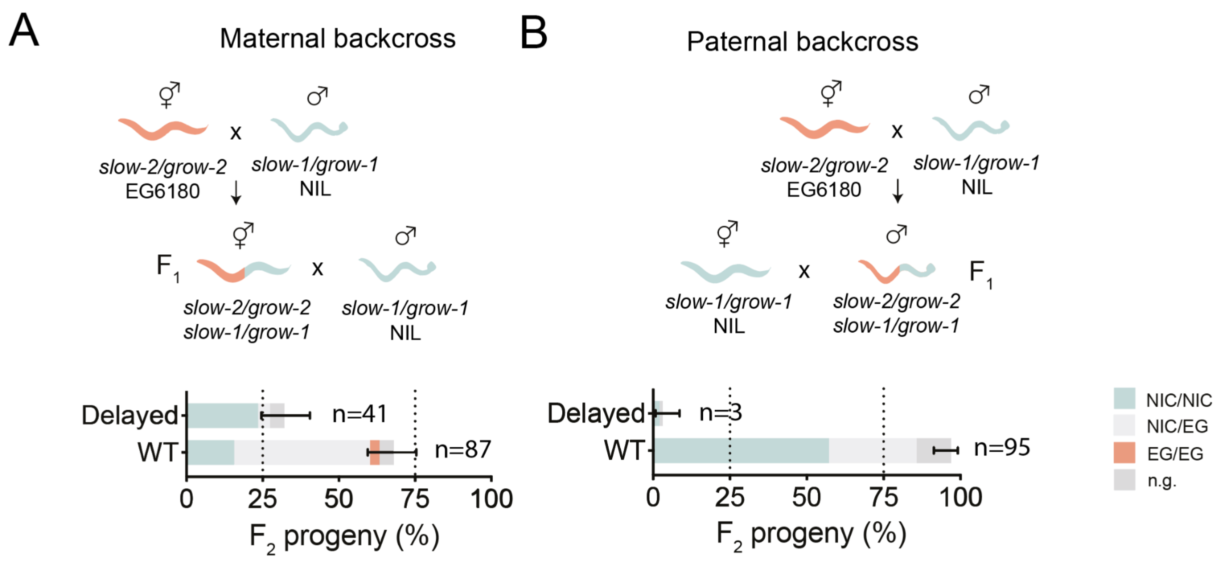


**Figure S6. *slow-2/grow-2* is a maternal-effect toxin-antidote element.** Reciprocal parental backcrosses **(A)** Maternal backcross of *slow-2/grow-2* TA. First, we crossed EG6180 hermaphrodites to *slow-1/grow-1* NIL males. We then crossed their heterozygous $F_1$ hermaphrodites (*slow-2/grow-2* / *slow-1/grow-1)* to parental *slow-1/grow-1* NIL males and inspected their $F_2$ progeny. **(B)** Paternal backcross of *slow-2/grow-2* TA. First, we crossed EG6180 hermaphrodites to *slow-1/grow-1* NIL males. We then crossed their heterozygous $F_1$ males (slow-2/grow-2 / *slow-1/grow-1)* to parental *slow-1/grow-1* NIL hermaphrodites and inspected their $F_2$ progeny. Delayed NIC/NIC individuals are observed when mothers carry the *slow-2/grow-2* TA but not males, thus showing that *slow-2/grow-2* is a maternal-effect TA.

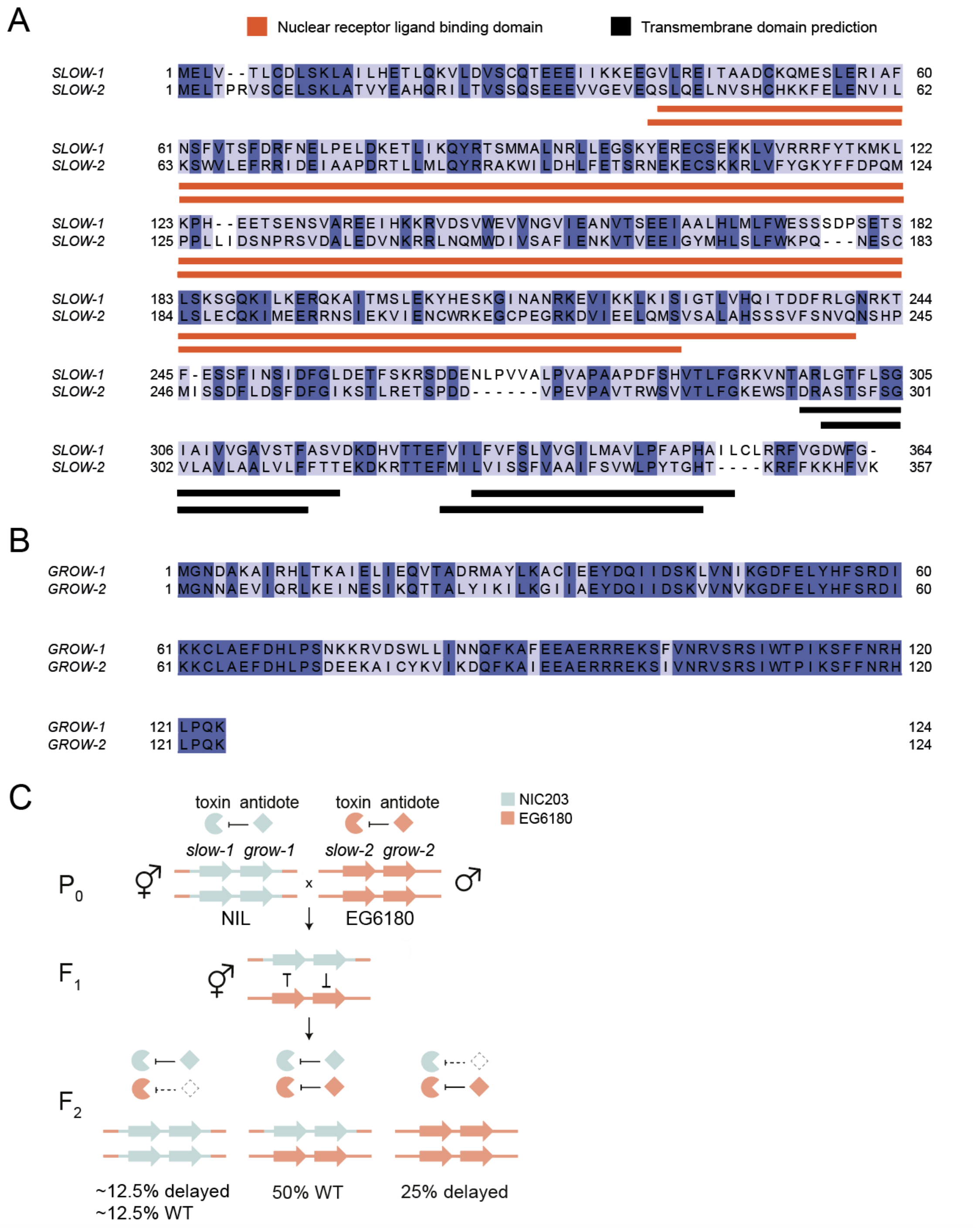


**Figure S7. Conservation and inheritance of *slow-1/grow-1* and *slow-2/grow-2.* (A)** Protein alignment of SLOW-1 and SLOW-2. Highlighted are predicted domains **(B)** Protein alignment of GROW-1 and GROW-2. The two proteins share 71% amino-acid pairwise identity with 88/124 identical sites. (B) Model illustrating the antagonism between the *slow-1/grow-1* and *slow-2/grow-2* TAs. Illustrated is a cross between Chr. III NIL hermaphrodites (*slow-1/grow-1* carriers) and EG6180 males (*slow-2/grow-2* carriers). Both TAs are found in the same locus and are mutually exclusive in NIC203 and EG6180 parental strains. $F_2$ progeny are poisoned by SLOW-1 and SLOW-2, two maternal-effect toxins. Heterozygous individuals are phenotypically WT because they express both zygotic antidotes. Homozygous *slow-2/grow-2* carriers are delayed because they lack the GROW-1 antidote and are poisoned by SLOW-1 (when maternally inheriteD). Homozygous *slow-1/grow-1* carriers are delayed because they lack the GROW-2 antidote and are poisoned by SLOW-2. Notice not all *slow-1/grow-1* carriers are affected, only about half of them. This could be due to some interference between the TAs or a lower activity of SLOW-2 compared to SLOW-1.

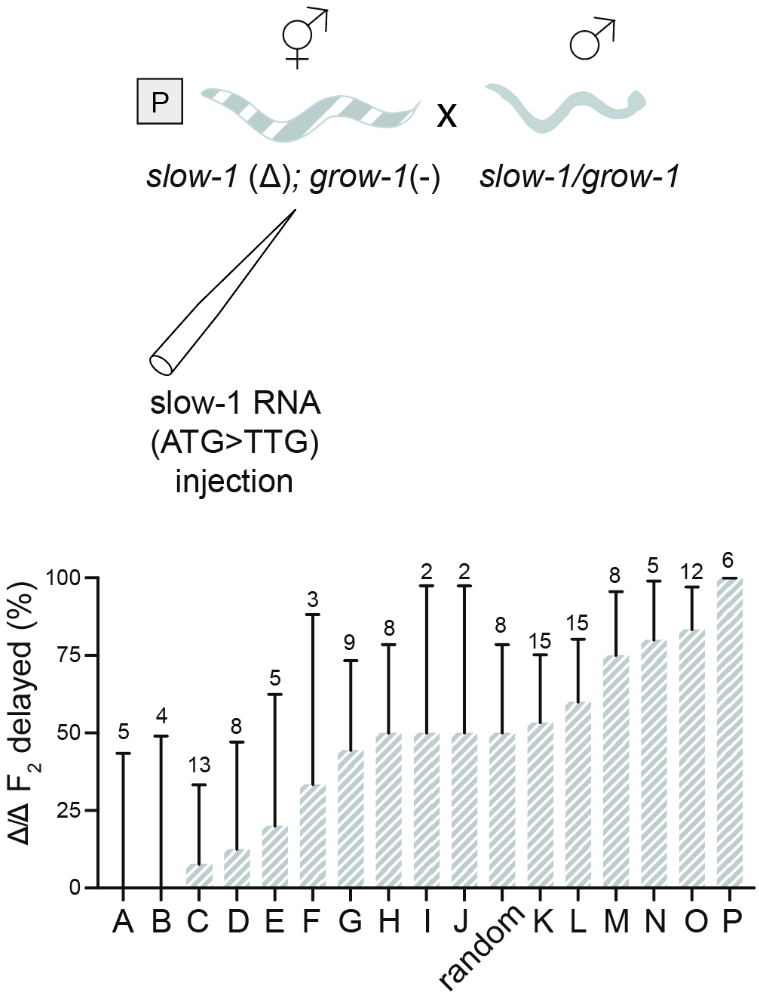


**Figure S8. Variability in the rescue of *slow-1/grow-1* activity across injected hermaphrodites following paternal inheritance.** Sixteen *slow-1(Δ)/grow-1*(-) NIL hermaphrodites (A to P) were injected in both gonad arms with i*n vitro* transcribed *slow-1* RNA (mutated start codon) and later mated to *slow-1/grow-1* NIL males. The total number of homozygous *slow-1(Δ)/grow-1*(-) $F_2$ progeny from each hermaphrodite is shown on top of each bar. The sample labeled as “random” represents embryos randomly picked from different injected mothers.

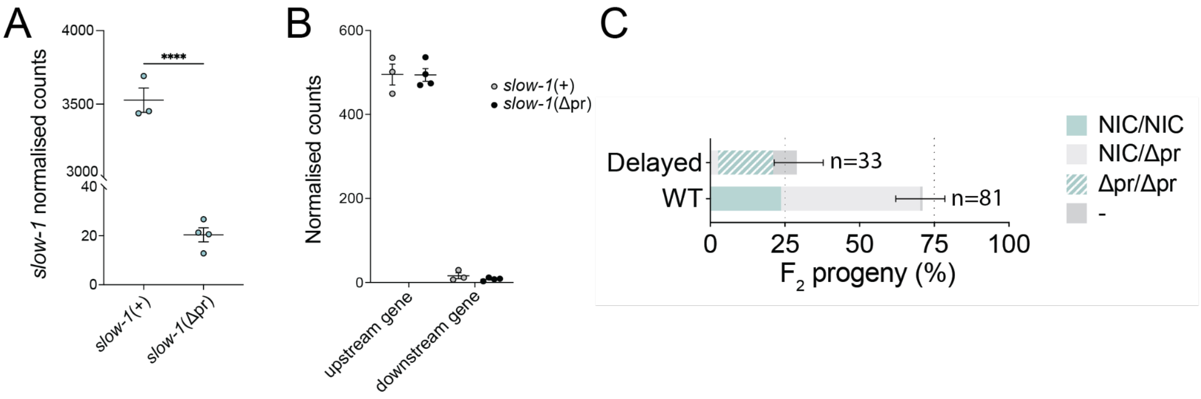


**Figure S9. A drastic decrease in maternal *slow-1* transcript abundance does not affect epigenetic licensing. (A)** Quantification of *slow-1* transcripts following deletion of a 620 bp region upstream of *slow-1*. Comparison between *slow-1*(+)/ *grow-1.1*(+)/ *grow-1.2*(-) NIL and *slow-1*(*Δ*pr)/*grow-1.1*(+)/ *grow-1.2*(-) NIL strains by RNA-seq. Deletion of the slow-1 promoter causes a 176-fold decrease in *slow-1* transcript levels (t-test, p<0.0001). **(B)** Quantification of the genes immediately upstream of *slow-1* and downstream of *grow-1*. Deletion of the *slow-1* promoter does not change the transcript level of the two neighboring genes (two-way ANOVA, interaction p=0.81). **(C)** Cross between *slow-1(Δpr)/grow-1*(-) NIL hermaphrodites and NIL males. All homozygous *slow-1(Δpr)/grow-1*(-) NIL $F_2$ progeny are delayed (100%, n=21).

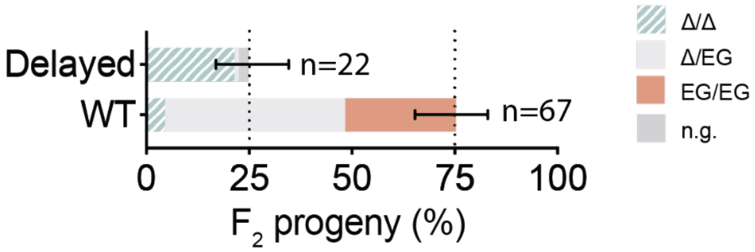


**Figure S10. Maternal *slow-1* mRNA does not license zygotic *slow-2*.** Phenotypes and genotypes of the $F_2$ progeny of a cross between *slow-1(Δ)/grow-1*(-) NIL hermaphrodites to EG6180 males. The results indicate that paternally-inherited *slow-2/grow-2* is still active in the absence of maternal *slow-1* mRNA.

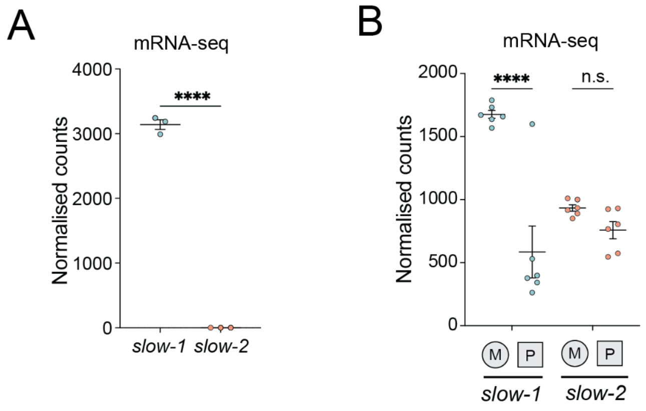


**Figure S11. The *slow-1* parent-of-origin effect is also present in a cross between NIC203 and EG62180 parental lines. (A)** Slow-1 transcripts are detected in NIC203 but not EG6180 by RNAseq (t-test, $p<0.0001$) **(B)** Reciprocal crosses between NIC203 and EG6180 parental strains. The abundance of *slow-1* transcripts is higher when the *slow-1* locus is maternally inherited (two-way ANOVA, interaction $p=0.0005$, Holm-Sidak post hoc test, $p_{slow\text{-}1}<0.0001$). In contrast, the transcript abundance of *slow-2* is similar when maternally and paternally inherited ($p_{slow\text{-}2}=0.27$).

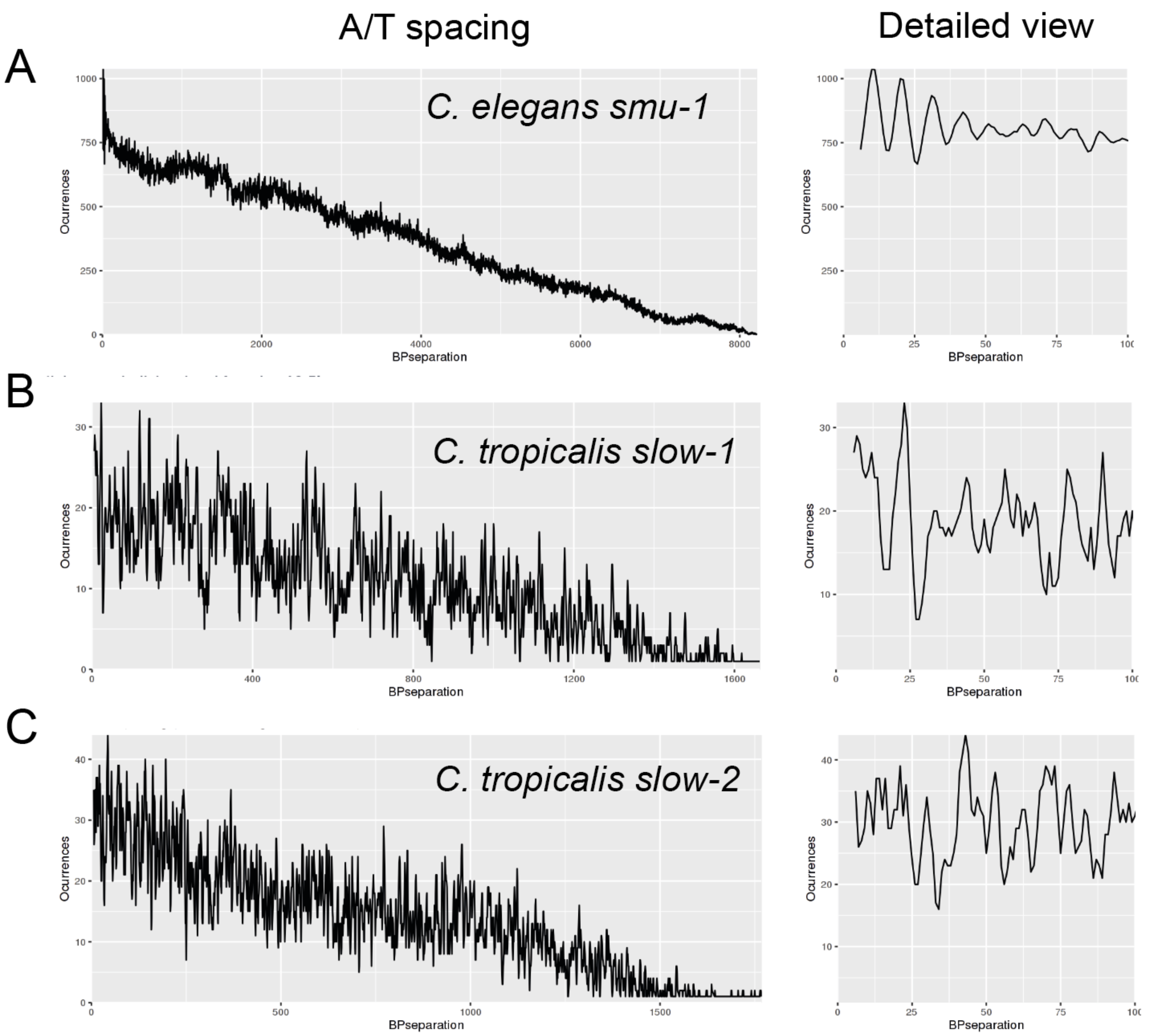


D

| | Sequence ID | Number of bases | Bases in phase | Phasing frequency | Total PATC value | Phasing density |
|---|---|---|---|---|---|---|
| 1 | smu-1 | 8250 | 3776 | 0.46 | 653206 | 79.18 |
| 2 | slow-1 | 1711 | 52 | 0.03 | 20083 | 11.74 |
| 3 | slow-2 | 1792 | 61 | 0.03 | 17698 | 9.88 |

**Figure S12. *Slow-1* and *slow-2* lack PATCs in their intronic sequences. (A)** PATC periodicity analysis for positive control *C. elegans smu-1*. Highest periodicity found at 10.5 bp. **(B)** PATC periodicity analysis for *C. tropicalis slow-1*. Highest periodicity found at 20 bp. **(C)** PATC periodicity analysis for *C. tropicalis slow-1*. Highest periodicity found at 14 bp. **(D)** Summary of PATC analysis. Phasing threshold was set to 60 (~ 1% Phasing in random DNA). Analysis was run on https://wormbuilder.org/patc/

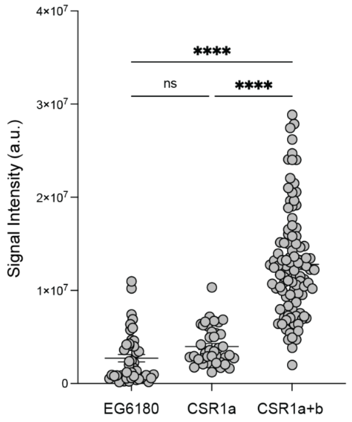


**Figure S13. Quantification of *C. tropicalis* CSR-1 expression at the 2-cell stage of embryonic development.** FLAG immunofluorescence quantification of 2-cell stage embryos of EG6180 (negative control), FLAG::CSR-1a and FLAG::CSR-1a+b (2 repeats, $n_{EG6180}$=44, $n_{CSR1a}$=40, $n_{CSR1a+b}$=105, one-way ANOVA, p<0.0001, Tukey post hoc test, $p_{EG\ vs\ CSR1a}$=0.41, $p_{EG\ vs\ CSR1a+b}$<0.0001, $p_{CSR1a\ vs\ CSR1a+b}$<0.0001).

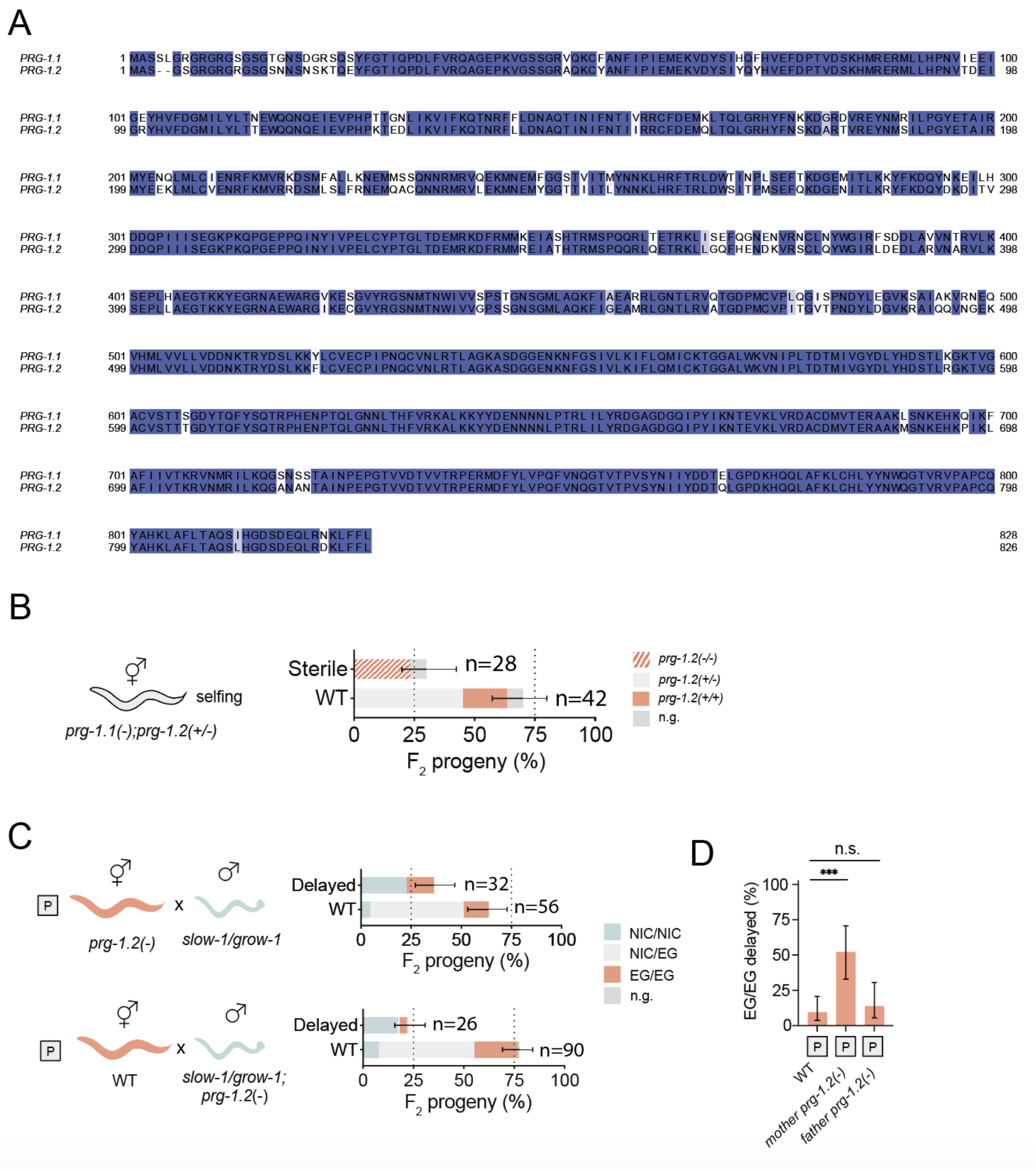


**Figure S14.** ***C. tropicalis*** **PRG-1.1 and PRG-1.2 are redundant paralogs and PRG.12 acts maternally (A)** Protein alignment of PRG-1.1 and PRG-1.2. The two proteins share 87% amino-acid pairwise identity with 722/828 identical sites. **(B)** Selfing of *prg-1.1*(-); *prg-1.2*(+/-) strain. All prg-*1.1*(-); *prg-1.2*(-) individuals were sterile, therefore the line couldn't be propagated. **(C)** Cross of *prg-1.2(-)* hermaphrodites to NIL males (top) and NIL hermaphrodites to *prg-1.2(-)* males (bottom) indicates that paternal slow-1 is repressed only when *prg-1.2* is maternally inherited. **(D)** Percentage of $F_2$ EG/EG delayed progeny when *prg-1.2* is absent from the mother or the father compared to the WT cross. ($n_p$=53, $n_{mother\ prg\text{-}1.2(-)}$=23, $n_{father\ prg\text{-}1.2(-)}$=29, Fisher's exact test compared to WT $p_{mother\ prg\text{-}1.2(-)}$=0.0001, $p_{father\ prg\text{-}1.2(-)}$=0.71).

### Supplementary Tables

| Strain | Short name | Species | Genotype | Description | Source |
|---|---|---|---|---|---|
| AF16 | AF16 | *C. briggsae* | wild type | Wild isolate from Ahmedabad, Gujarat, India. Collected by A. Fodor | CGC |
| DL238 | DL238 | *C. elegans* | wild type | Wild isolate from Manuka National Reserve, Hawaii, USA | CGC |
| EG6180 | EG6180 | *C. tropicalis* | wild type | Wild isolate from El Yunque, Puerto Rico. Lat 18.3 Lon -65.8. Found in rotting fruit by M. Ailion and E. Jorgensen | Christian Braendle |
| HK104 | HK104 | *C. briggsae* | wild type | Wild isolate from Okayama, Japan. Collected by S. Baird | CGC |
| INK112 | *slow-2(-)/ grow-2(-)* | *C. tropicalis* | *slow-2(abu70*[p.C71X]*)*; *grow-2(abu58*[p.7VX]*)* III; EG6180 | *slow-2 grow-2* double mutant in EG6180 background | This study |
| INK124 | *slow-2(-)* | *C. tropicalis* | *slow-2(abu70*[p.C71X]*)* III; EG6180 | *slow-2* mutant in EG6180 background | This study |
| INK255 | Chr. III NIL *dpy* | *C. tropicalis* | *dpy(abu148)* X; *qqIR46* | dpy mutant in QX2345 NIL background | This study |
| INK264 | *prg-1.2(-)* | *C. tropicalis* | *prg-1.2(abu185*[Y183X]*)* I; EG6180 | *prg1.2* mutant in EG6180 background | This study |
| INK266 | *prg-1.1(-)* | *C. tropicalis* | *prg-1.1(abu183*[M189X]*)* I; EG6180 | *prg1.1* mutant in EG6180 background | This study |
| INK293 | slow-1(Δ)/ grow-1(-) Chr. III NIL | *C. tropicalis* | *slow-1(abu122*[Δ*slow-1*]*)*; *grow-1.1(abu178*[p.T12SfsX2]*)*; *grow-1.2(abu132*[p.T12SfsX2]*)* III; qqIR46 | *slow-1* full coding deletion and grow-1.1 grow-1.2 mutant in QX2345 NIL background | This study |
| INK304 | *prg-1.1(-)* Chr. III NIL | *C. tropicalis* | *prg-1.1(abu173*[M189X]*)* I; *qqIR46* | *prg1.1* mutant in QX2345 NIL background | This study |
| INK305 | *prg-1.2*(-) Chr. III NIL | *C. tropicalis* | *prg-1.2(abu185*[Y183X]*)* I; *qqIR46* | *prg1.2* mutant in QX2345 NIL background | This study |
| INK348 | FLAG:: CSR-1a | *C. tropicalis* | *csr-1*(abu235[N1-3xFLAG::*csr-1*]) *IV* | FLAG::CSR-1a tagged strain | This study |
| INK351 | *myo-2p:: mScarlet* EG6180 | *C. tropicalis* | *abuSi10[myo-2p::mScarlet::unc-54 3'UTR + HygR(+); abuSi9] IV* | EG6180 with red fluoresecent pharynx. Single copy insert in Chr. IV abuSi9 landing site. Transgene used as a visual marker for crosses | This study |
| INK364 | *csr-1*::Neo/+ | *C. tropicalis* | *csr-1(abu239*[*csr-1::p.Leu337X::rps-20p::NeoR::rps-20* 3'UTR]) IV/+ | Balanced csr-1 null strain. Propagate on G418. Only hets and a few csr-1(-) survive | This study |
| INK367 | *csr*-1(-) | *C. tropicalis* | *csr-1(abu239*[*csr-1::p.Leu337X::rps-20p::NeoR::rps-20* 3'UTR]) IV/+; *qqIR46* | *csr-1* null strain. High embryonic and larval lethality but viable | This study |
| INK372 | csr-1::Neo/+ Chr. III NIL | *C. tropicalis* | *csr-1(abu239*[*csr-1::p.Leu337X::rps-20p::NeoR::rps-20* 3'UTR]) IV/+; *qqIR46* | Balanced csr-1 null strain in QX2345 NIL background. Propagate on G418. Only hets and a few csr-1(-) survive | This study |

| | | | | | |
|---|---|---|---|---|---|
| INK374 | PRG-1.1:: FLAG | *C. tropicalis* | *prg-1.1(abu250*[3xFLAG::*prg1.1*]) I | PRG-1.1 FLAG tagged strain in EG6180 background | This study |
| INK377 | PRG-1.2:: FLAG | *C. tropicalis* | *prg-1.2(abu253*[3xFLAG::*prg1.2*]) I | PRG-1.2 FLAG tagged strain in EG6180 background | This study |
| INK387 | FLAG:: CSR-1a+b | *C. tropicalis* | *csr-1(abu258*[N2-FLAG::*csr-1*]*)* IV | FLAG::CSR-1a+b tagged strain | This study |
| INK442 | *slow-1(Δpr)/grow-1(-)* Chr. III NIL | *C. tropicalis* | *slow-1(abu313*[Δprom::*slow-1*]*); grow-1.1(abu310*[p.T12SfsX2]*); grow-1.2(abu154*[p.T12SfsX2]*)* III; *qqIR46* | *slow-1* promoter deletion and grow-1.1 grow-1.2 mutant in QX2345 NIL background | This study |
| INK459 | mScarlet:: SLOW-1 | *C. tropicalis* | *slow-1(abu287*[*mScarlet::slow-1*]*)* III; *qqIR46* | mScarlet N-terminal endogenously tagged *slow-1* in NIL background | This study |
| INK460 | *Myo-2p:: mScarlet* Chr. III NIL | *C. tropicalis* | *abuSi10[myo-2p::mScarlet::unc-54 3'UTR + HygR(+); abuSi9] IV; qqIR46* | Chr. III NIL with red fluoresecent pharynx. Single copy insert in Chr. IV abuSi9 landing site. Transgene used as a visual marker for crosses | This study |
| INK461 | mScarlet:: SLOW-1 *dpy* | *C. tropicalis* | *slow-1(abu287*[*mScarlet::slow-1*]*)* III; *dpy(abu148)* X; *qqIR46* | mScarlet N-terminal endogenously tagged *slow-1* in NIL background and recessive dpy mutation in Chr. X | This study |
| INK531 | NIC203 *dpy* | *C. tropicalis* | *unc(abu308) X; NIC203* | Uncoordinated mutant in NIC203 background obtained by EMS mutagenesis. Recessive allele. Backcrossed 4X into NIC203. Mutation maps to Chr. X | This study |
| N2 | N2 | *C. elegans* | wild type | Reference strain | CGC |
| NIC203 | NIC203 | *C. tropicalis* | wild type | Wild isolate from Capesterre Belle-Eau, Guadeloupe. Lat 16.05 Lon -61.63. Found in rotting flowers by N. Poullet and C. Braendle | Christian Braendle |
| QX2341 | Chr II NIL | *C. tropicalis* | *qqIR45 (II:8.0-8.7 Mb; NIC203 > EG6180); EG6180 Mito* | NIL carrying NIC203 TA element Chr. II in a EG6180 background | (20) |
| QX2345 | Chr. III NIL | *C. tropicalis* | *qqIR46 (III:10.42-10.46 Mb; NIC203 > EG6180)* | NIL carrying NIC203 TA element Chr. III in a EG6180 background | (20) |
| QX2355 | EG6180 dpy | *C. tropicalis* | *dpy(qq104) X;* EG6180 | Spontaneous dumpy recessive mutation in EG6180 background. Mutation maps to Chr. X | (20) |
| QX2361 | *slow-1(-)/ grow-1(-) NIL* | *C. tropicalis* | *slow-1 (qq100[p.H125RfsX15]); grow-1.1(qq101[p.T12SfsX2]); grow-1.2(qq101[p.T12SfsX2]) III; qqIR46* | *slow-1 grow-1.1 grow1.2 mutant in QX2345 NIL background* | (20) |

### Table S1. List of strains used in the study

| Marker/ Gene | FW primer | RV primer | Sequencing primer/gel | Comments |
|---|---|---|---|---|
| NIC203 Chr. III TA | GCCTAGAAAAACAATTGATGGCC | CGAGTAATTTCACCGGGTTG | gel | Deletion in EG6180. 699 bp in NIC203 and 556 bp in EG6180 |
| *grow-1.1* frameshift | AAATAGGCGGGGCTTTCTTA | GGAGGCAGGAGAGTCCTTCT | ACCGGTAAATGGTCGAATTCAGC | 1125 bp |
| *grow-1.2* frameshift | TGCTGAGGAGTCTTCGATTCCC | ATCTCATTTTCCCGCCAAACGC | ACCGGTAAATGGTCGAATTCAGC | 1085 bp |
| *slow-2* frameshift | AAGCCAATATGGAGTTGACGCC | TAACGGAGGCATCTGTGGATCG | GCGTATCATGCGAACTCTCCAA | 458 bp |
| *grow-2* frameshift | TCTCGAGAATTTCTGCACTGTTCAA | TGTACTGCATCCTCCGACGTTT | TCTCGAGAATTTCTGCACTGTTCAA | 655 bp |

**Table S2. Primers used for genotyping of the genetic crosses**

| Gene | Modification | Sequence |
|---|---|---|
| *slow-1* | *slow-1* deletion | ATTTTCCTGGGAATCAACAC |
| *slow-1* | *slow-1* deletion | TCCGTGTCAGCACAATATAT |
| *slow-1* | *slow-1* promoter deletion | AAATCCTCATTTCCCGTCAA |
| *slow-1* | *slow-1* promoter deletion | CGTGAATGCAGGAAACTCGC |
| *slow-1* | *slow-1* promoter deletion | AAAACCATTGACGGGAAATG |
| *slow-1* | *slow-1* promoter deletion | AAGTGTAGAATTTGCAGAAA |
| *slow-1* | mScarlet::*slow-1* | AGAGATCACAGAGCGTTACAA |
| *slow-2* | *slow-2* frameshift | TTGTCACAAGAAATTCGAGT |
| *grow-2* | *grow-2* 3xStop | TATTCATTGATTCTATGCAA |
| *prg-1.1* | *prg-1.1* frameshift | TATAACATGAGAATTCTCCC |
| *prg-1.2* | *prg-1.2* frameshift | CTTTTGAGTTGAAGTAGTGA |
| *prg-1.1* | FLAG::*prg-1.1* | GTAAATATGGCTTCCAGTTT |
| *prg-1.2* | FLAG::*prg-1.2* | TACCGGATGCCATTATTACC |
| *csr-1* | neoR insertion | AAGAACGGTTTATCATGATC |
| *csr-1* | neoR insertion | CATCTCAAGGAGCAATCAGA |
| *csr-1* | FLAG::*csr-1a+b* | AACCGTGGACGAGATACTAG |
| *csr-1* | FLAG::*csr-1a* | CCGTTCGAGTTCATTTTTGA |

**Table S3. gRNAs used in this study**

| Gene | Modification | Sequence |
|---|---|---|
| *slow-1* | *slow-1* deletion | CTATCTTCAAGGGGTCGGGGCCTAGGAAAAGTGGGCGGAGTTTGAAATTTAAATAGGCGGGGCTTGTGG<br>AGAAAGTGGGCGTGGCCAGTGTATTGGCGGTAATTCAAATTCCGTTTCTTTTGTTTCCCATTTTTCTTG<br>ATTTTTCTCCGACAAAAATTACTTTTTTGAGTCAGAAATGAT |
| *slow-1* | *slow-1* promoter deletion | GCGAATTTCTTGAGTTTTCAAGTGATTTAGAAACAGAAACACGTAGAAAAGTGTAGAATTTGCAGAAAC<br>CAGATGAAATGGAGCTTGTAACGCTCTGTGATCTCTCAAAGCTTGCAATTCTCCACGAAACACTGC |
| *grow-2* | *grow-2* 3xStop | CTCTCTCTCCCTCACTCATCTAGTCCCCTCCTTGCGCGTATTCATTGATTCTATGCAATGGGAAACAAC<br>GCCGAATGACTAATTGAAAGAAATTAACGAATCAATAAAACAGACAACGGTGATCTTTCAGAAATTCCT<br>AATA |
| *prg-1.1* | FLAG::*prg-1.1* | GGACTTTATTTCTTTGTAATTATCAACTTAATTACATCAATAATTTTTTCAGGTAAATATGGATTACAA<br>AGACCATGATGGTGACTATAAGGATCATGATATTGACTATAAAGATGACGATGACAAGGCTAGTTCCTT<br>AGGTAGAGGCAGAGGACGCGGCTCTGGGTCAGGAACTGGAAACAGTGATGG |
| *prg-1.2* | FLAG::*prg-1.2* | GGAGTTGCTGTTGTTAGATCCAGATCCACGTCCTCTGCCTCTTCCACTACCGGATGCCTTGTCATCGTC<br>ATCTTTATAGTCAATATCATGATCCTTATAGTCACCATCATGGTCTTTGTAATCCATTATTACCTGGAA<br>AAAATATGCGAAAAATCTAGGAGAAACCGAGGAGAATCGACGG |
| *csr-1* | FLAG::*csr-1a+b* | GCGGGTTTCGAGAAAAGTAACAATTTTCAGCACAAAATGCAGTCTGGGAATTCTAACCGTGGACGAGAT<br>TACAAAGACCATGATGGTGACTATAAGGATCATGATATTGACTATAAAGATGACGATGACAAGACTAGG<br>GGGAATGATCGTGGAAATAGTGGAAGAGGTGGACGTGGATCGACTAGAGGTAAAAGAGG |
| *csr-1* | FLAG::*csr-1a* | GCATCTTCACACCTAGCTTAACTTCAGATAATCCGTCAAAAATGGATTACAAAGACCATGATGGAGACT<br>ATAAGGATCATGATATTGACTATAAAGATGACGATGACAAGAACAGTAATGGTAACCCCAGACTGGCAA<br>TTAACATTTTTGGACTTGAGCTTTCCGAACGCAAGATTTTCCG |

**Table S4. Homology-Directed Repair templates used in this study**

| 3’ adaptor | /5rApp/NN NNN NXX XXX AGA TCG GAA GAG CAC ACG TCT /3ddC/ |
|---|---|
| sRBC-1001 | CAGTG |
| sRBC-1002 | AGCAA |
| sRBC-1003 | GGTAT |
| sRBC-1004 | TACCA |
| sRBC-1005 | GTCAG |
| sRBC-1006 | TGACT |
| sRBC-1008 | CGTTC |
| sRBC-1009 | ATGGA |
| sRBC-1010 | GAACG |
| sRBC-1011 | ACGAG |
| 5’ adaptor | ACACUCUUUCCCUACACGACGCUCUUCCGAUCUNNNNN |

**Table S5. sRNA library adaptors and sRNA barcodes (sRBC, indicated as X)**
